# A rule-based data-informed cellular consensus map of the human mononuclear phagocyte cell space

**DOI:** 10.1101/658179

**Authors:** Patrick Günther, Branko Cirovic, Kevin Baßler, Kristian Händler, Matthias Becker, Charles Antoine Dutertre, Venetia Bigley, Evan Newell, Matthew Collin, Florent Ginhoux, Andreas Schlitzer, Joachim L. Schultze

## Abstract

Single-cell genomic techniques are opening new avenues to understand the basic units of life. Large international efforts, such as those to derive a Human Cell Atlas, are driving progress in this area; here, cellular map generation is key. To expedite the inevitable iterations of these underlying maps, we have developed a rule-based data-informed approach to build next generation cellular consensus maps. Using the human dendritic-cell and monocyte compartment in peripheral blood as an example, we performed computational integration of previous, partially overlapping maps using an approach we termed ‘backmapping’, combined with multi-color flow-cytometry and index sorting-based single-cell RNA-sequencing. Our general strategy can be applied to any atlas generation for humans and other species.

**Graphical Abstract:** 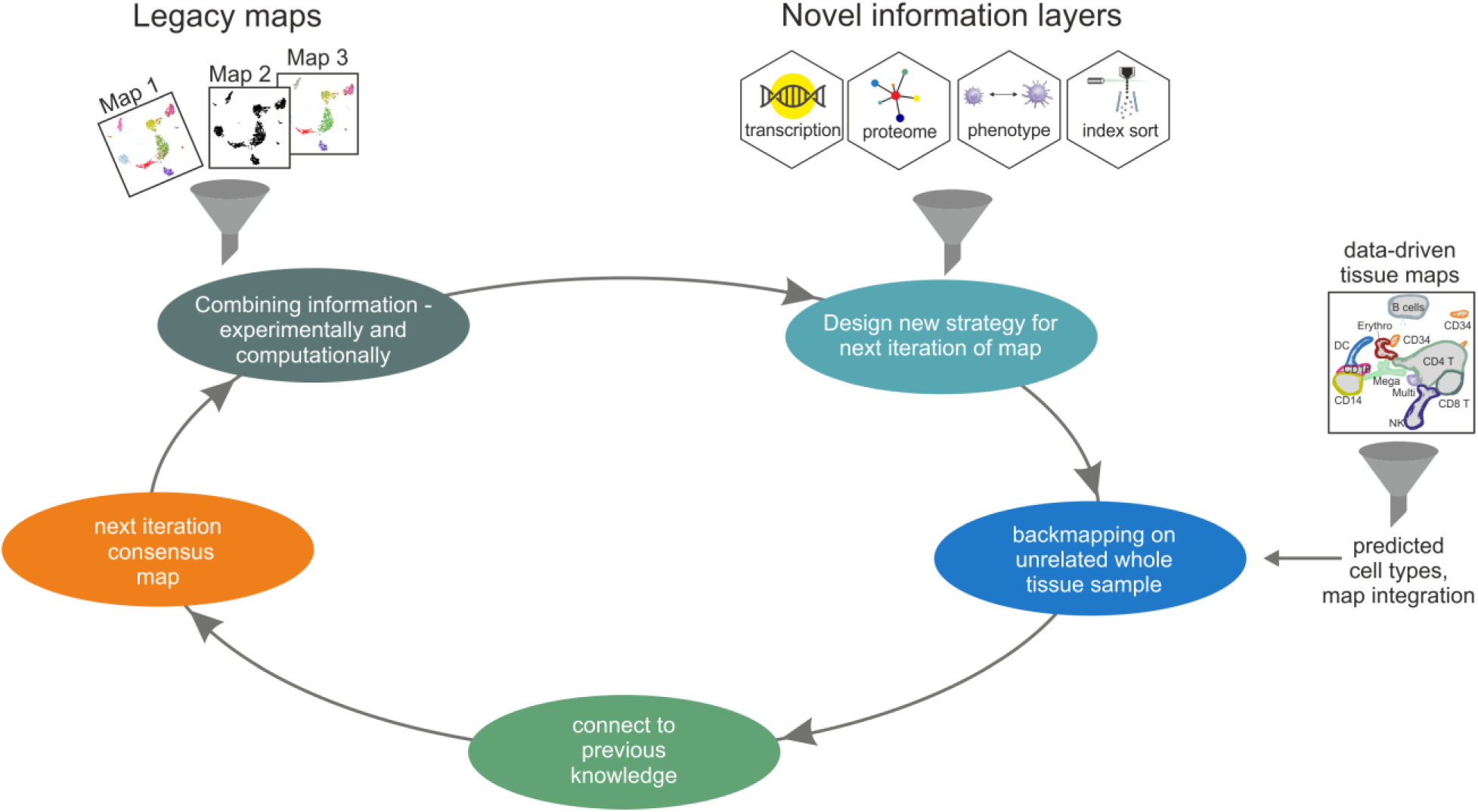

**Highlights:**

- Defining a consensus of the human myeloid cell compartment in peripheral blood
- 3 monocytes subsets, pDC, cDC1, DC2, DC3 and precursor DC make up the compartment
- Distinguish myeloid cell compartment from other cell spaces, e.g. the NK cell space
- Providing a generalizable method for building consensus maps for the life sciences

## Introduction

Since Robert Hooke’s first observations of cells as the basic unit of life, generations of life scientists have been driven to understand, map and characterize individual cells (Cavaillon, 2011). For many decades, morphological parameters were the major driving force to establish new cell identities (Hussein et al., 2015). In immunology, technologies such as flow cytometry have been developed that permit quantitative enumeration of single cells based on measuring combinations of predominantly cell-surface proteins (Hulett et al., 1969; Perfetto et al., 2004). These technologies, however, have some undisputable limitations, most notably, their reliance on a predefined subset of biomolecules. Conversely, single-cell-omics, particularly single-cell transcriptomics, allow for cells to be assessed, in principle, without predefined markers. Here, the complete spectrum of transcriptomic parameters is investigated and used as a defining unit of cell identity (Islam et al., 2014; Macosko et al., 2015; Tang et al., 2009). Such single-cell technologies allow for a fully data-driven analysis to establish cell maps of an organism, such as those proposed by the Human Cell Atlas consortium (Rozenblatt-Rosen et al., 2017). We have learnt from other disciplines that maps require iterations over time, often due to new data generated as a result of technological advances. These iterations improve the precision, accuracy and available content per data point (Edney, 2019; Monmonier, 2015; Ridpath, 2007).

Reliable consensus maps are a prerequisite to reconcile conflicting data that might have been generated based on different data generating approaches (Edney, 2019; Monmonier, 2015). Here we generalize the approach of building geographic or astronomic consensus maps to human cellular consensus maps. We exemplify our approach by integrating two recently introduced single-cell transcriptomics-based cellular maps of the human blood mononuclear myeloid cell compartment (See et al., 2017; Villani et al., 2017) with novel single-cell transcriptomics and flow cytometry data. The human blood mononuclear myeloid cell compartment has been recognized to harbor a complex mixture of cells of diverse origins exemplified by the ongoing efforts to map this cellular compartment with increasing resolution (Dutertre et al., 2019). The two mapping efforts present with discrepancies and commonalties in terms of cell type identification, naming and breadth of sampling. In order to establish a consensus map of the human mononuclear myeloid cell compartment we allow for the integration of prior knowledge in that we define *a priori* criteria for the cellular compartment under study in order to increase resolution and to allow building of a consensus map. Overall, our approach generates rule-based data-informed cellular consensus maps that resolve discrepancies between the two recently generated maps, and clarifies cellular identities of human dendritic-cell (DC) and monocyte subsets resulting in a novel, integrated consensus map of the human blood myeloid compartment.

## Results

### Integrated phenotypic characterization of the myeloid cell compartment in human peripheral blood

We aimed to build a consensus map of healthy human blood myeloid cells that integrates legacy dataset knowledge into a revised consensus map. To do so, we generated a novel single-cell-omics dataset of the blood CD45^+^Lin^-^HLA-DR^+^ cell space using a 17 parameter index sorting panel incorporating important markers from two recently published single-cell–omics datasets, here termed map 1 (Villani et al., 2017) and 2 (See et al., 2017) and an established panel of myeloid cell markers including CD14, CD16, HLA-DR, CD1c and CADM1 (Dutertre et al., 2014; Guilliams et al., 2016; Haniffa et al., 2012), to link the data to the body of knowledge already present within the literature (Figure 1A, S1A-C, Table S1). This strategy allowed us to directly include several cell populations defined by either map 1 or 2 into our single-cell transcriptomics dataset, compare these populations within an unbiased myeloid cell space dataset, and assess differences and commonalities between the two maps.

**Figure 1.**
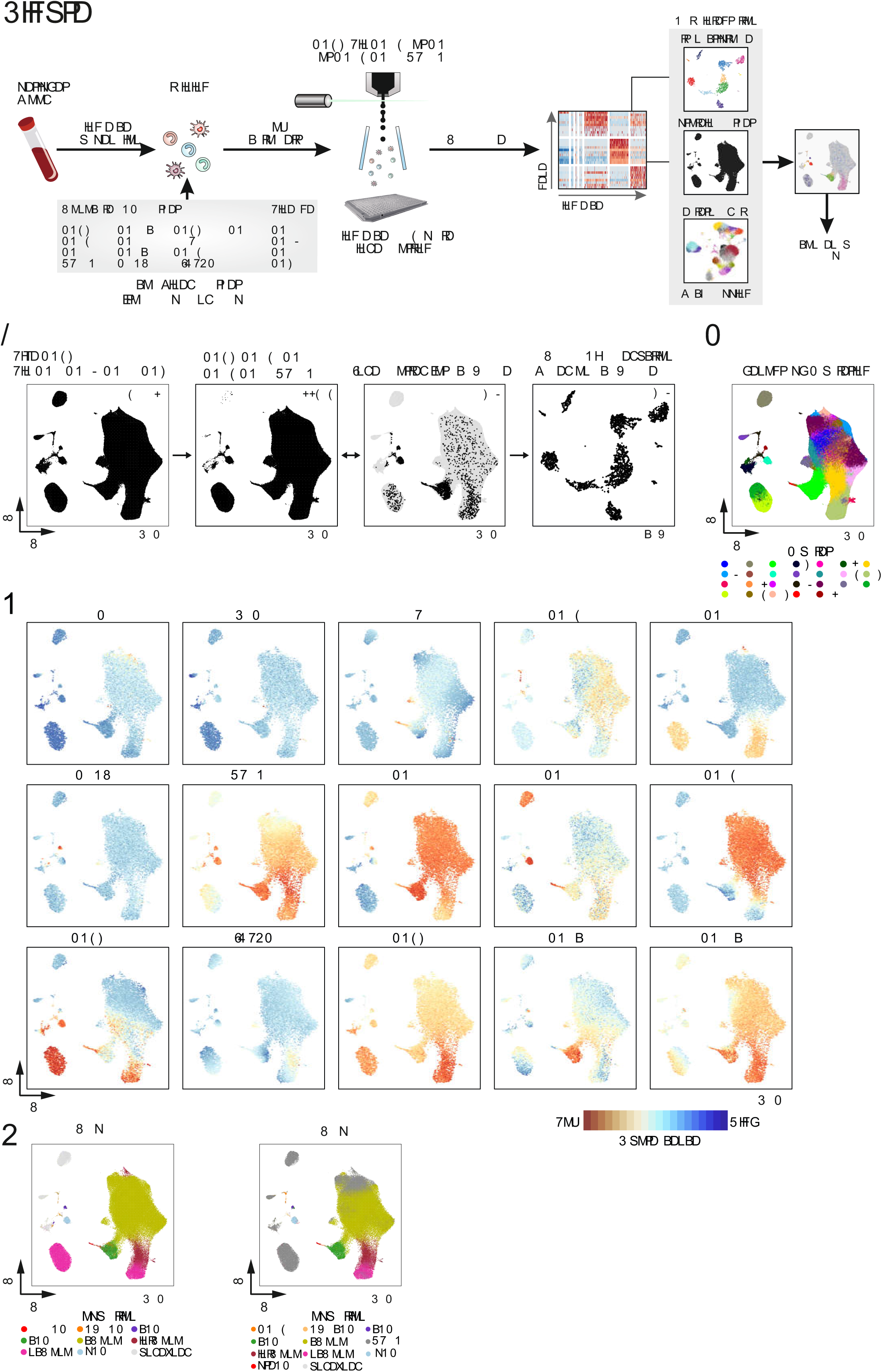
Generating a new consensus map of the mononuclear myeloid cell compartment in human peripheral blood. **(A)** Workflow to generate a new consensus map of the human mononuclear myeloid cell compartment. **(B)** Visualization of ∼1.4 mio. live CD45^+^Lin(CD3, CD19, CD20, CD56)^-^ cells after UMAP dimensionality reduction of the flow cytometry panel introduced in A (left panel), mononuclear myeloid cell compartment (second panel), overlay of index-sorted cells (third panel), UMAP topology of the index-sorted cells based on the single-cell transcriptome data (most right panel, see also Figure 2). Grey areas in the third panel represent the CD45^+^Lin^-^ cell space. **(C)** Phenograph clustering of the flow cytometry data projected onto the FACS-based UMAP topology. **(D)** Color-coded visualization of markers used to define the mononuclear myeloid cell compartment. **(E)** Overlay of the cell gating strategies according to maps 1 (Villani et al., 2017) and 2 (See et al., 2017). See also Figure S1.

To understand the organization of the blood-derived myeloid cell compartment, we performed dimensionality reduction using the uniform manifold approximation and projection (UMAP) algorithm (Becht et al., 2018) on the complete flow cytometry space of live CD45^+^ Lin^-^ cells (Figure 1B). UMAP revealed a complex topology of the flow cytometry data, segregating a large cluster on the right and multiple small entities on the left of the topology. A fraction of the Lin^-^ cells (Figure 1C, cluster two) was not part of the monocyte or DC cell space according to CD16, CD14 and HLA-DR expression (Figure 1D). These cells most likely represent basophils due to their lack of HLA-DR expression but high CD123 expression (Figure 1D, Figure S1). To fully understand the population structure of the presented FACS-based UMAP, we performed Phenograph clustering of the live CD45^+^ Lin^-^ blood-derived flow cytometry UMAP space and detected 27 clusters (Figure 1C). To link these novel data to the two existing maps for the blood myeloid cell compartment (Guilliams et al., 2014; See et al., 2017; Villani et al., 2017) we reapplied the gating strategies of either map 1 or map 2 and overlaid these onto our novel flow cytometry-derived UMAP topology (Figure 1E, Figure S1A-C). This analysis revealed several commonalities and discrepancies between maps 1 and 2 in the combined novel flow cytometry panel used in this study. On the upper-most level, map 1 was less stringent in excluding HLA-DR^-^ cells within the myeloid cell space (cells labeled light grey, Figure 1D, S1B), a feature rigorously adhered to in map 2 (cells labeled dark grey, Figure 1D, S1C). Furthermore, Axl^+^Siglec6^+^ DCs (AS-DC; DC 5, Table S1) in map 1 occupied the same topological space as pre-DCs in map 2, indicating potential cellular overlap. Finally, map 1 mono 2/4, resembling non-classical monocytes (ncMono) (Table S1), occupied two different locations on the UMAP topology: one of them being within the HLA-DR^-^ compartment of the topology and the other being within the space assigned to monocytes by a classical investigator-derived flow cytometry gating (Figure 1D, 1E, S1). These data suggest that there is a commonality in the identity of map 1 Axl^+^Siglec6^+^ DCs (AS-DC; DC 5) and map 2 pre-DCs whereas mono 2/4 may represent a heterogeneous mixture of various cell types – apparently not all of them related to the myeloid cell lineage.

### Novel integrated single cell-omics data identifies commonalties and discrepancies between two recent myeloid cell maps

To investigate the cell population structure at the transcriptomic level we performed single-cell RNA-sequencing (scRNA-seq) of 2,509 blood-derived single cells following index sorting to encompass all major populations identified in either map 1 or 2 after lineage exclusion and generated a UMAP dimensionality reduction-based transcriptome map (Figure 1B, S2A-F). *De novo* clustering of the scRNA-seq data revealed 11 transcriptionally different clusters (Figure 2A, 2B, **Data Table S1**). We projected the cluster identities onto the flow cytometry-derived UMAP topology, which allowed us to validate our index sorting strategy and link identities across the flow cytometry and scRNA-seq data (Figure 2A).

**Figure 2.**
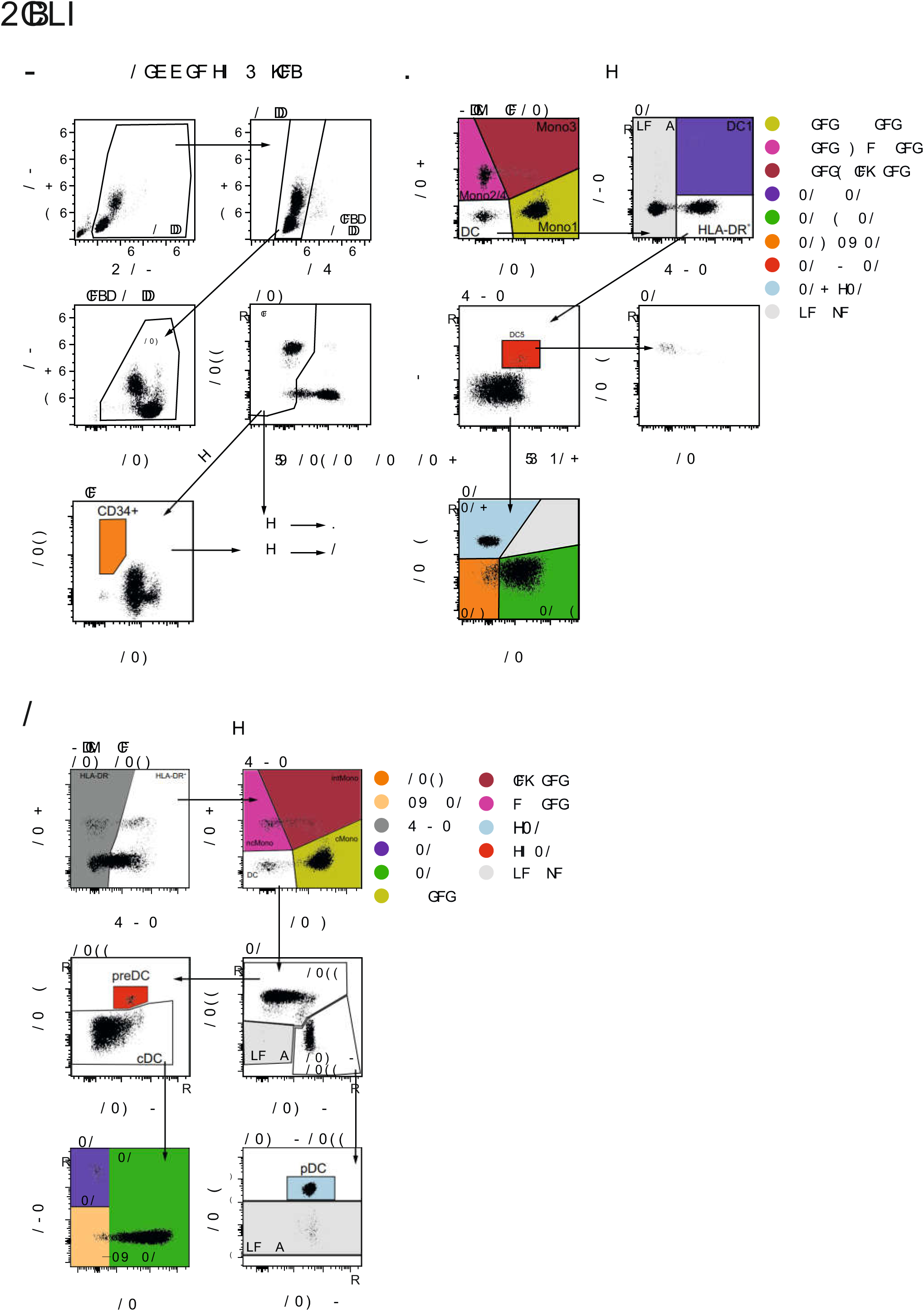
Index-sorted scRNA-seq dataset of the myeloid cell compartment in human blood. **(A)** *De novo* clustering of the 2,509 index-sorted cells onto the scRNA-based UMAP topology (left panel) and cluster projection onto the FACS-based UMAP topology (right panel, grey background: complete CD45^+^Lin^-^ cell space). **(B)** Heatmap of 10 most significant marker genes for each of the 11 clusters identified and visualized in Figure 2A. **(C)** Overlay of cell types defined for maps 1 (left panel) and 2 (right panel) onto the scRNA-based UMAP topology of the new consensus map.

Gene level inspection of these clusters revealed that cluster one had a natural killer (NK) cell signature, as indicated by PRF1, GNLY and NKG7 gene expression. Cluster two was identified by high IGH family gene expression, thus implying contaminating B cells with a strong plasmablast signature (Shi et al., 2015) (Figure 2B). Cluster three was represented by a minor fraction of the cells within our dataset, with a profile of microRNA-related transcripts. Cluster four expressed SPINK2, GAS5, SATB1and STMN1 genes, and thus corresponded to blood-derived CD34^+^ hematopoietic stem cells (Satoh et al., 2013; Will et al., 2013). Cluster five expressed the plasmacytoid DC (pDC)-related IRF7, TCF4 and GZMB transcripts (See et al., 2017; Villani et al., 2017), whereas cluster six expressed a conventional dendritic cell 1 (cDC1) gene-set, with high expression of CLEC9A, IDO1 and CD74 (van der Aa et al., 2015; Zhang et al., 2012). Interestingly, cluster seven expressed genes either affiliated to pre-DCs (See et al., 2017) or AS-DCs (DC5) (Villani et al., 2017), such as SIGLEC6, AXL, PLAC8 or LILRA4, thus associating them to the human pre-DC continuum. As expected from our sorting strategy, we also detected several clusters belonging to the monocyte lineage. Cluster eight represented CD16^+^ ncMono cells based on high FCGR3A (CD16) with SERPINA1 and DUSP6 expression. Conversely, cluster nine expressed S100A8, S100A9 and S100A12 together with VCAN and FCN1, identifying them as classical CD14^+^ monocytes (Mono1/cMono). Clusters 10 and 11 represented two cDC2 identities (DC2, DC3): both clusters expressed high levels of the cDC2-related CD1C, CD1E and several HLA-DR transcripts. Interestingly, and as shown in map 1 (Villani et al., 2017), cluster 11 co-expressed certain monocyte-affiliated gene products, such as S100A8, S100A89 and FCN1 (Figure 2B), as also shown in Dutertre et al. (Dutertre et al., 2019).

To develop our consensus map, we utilized the index sorting data of the populations identified in map 1 and 2 and mapped them onto our single-cell transcriptomic dataset (Figure 2C). Overlaying this index-sorting data onto the scRNA data-derived UMAP topology reiterated several commonalities between maps 1 and 2, including DC1/cDC1 (purple), DC6/pDC (pink), Mono1/cMono (ochre) and CD14^+^CD16^+^ intermediate monocytes (Mono3/intMono, dark red). Importantly, the detected discrepancies between maps 1 and 2 were also apparent on the transcriptomic level. Firstly, we noticed that map 1 double negative DCs (DN-DC) populated the same position within the UMAP topology as cDC2s in map 2. Furthermore, mapping the index sorting data of the map 1 ncMono population (transcriptionally defined in map 1 as Mono2 and 4, Table S1) revealed two separate cell clusters within the transcriptomic UMAP topology, indicating considerable cell-type heterogeneity within this population as defined by map 1. Interestingly one of the Mono2/4 clusters overlapped with the ncMono (magenta) cluster, whereas another cluster was mapped as HLA-DR^-^ within the flow cytometry gating strategy used in map 2 (Figure S1). Moreover, we noticed that map 1 AS-DCs (red) and map 2 pre-DCs (red) occupied the same topological space, indicating considerable transcriptomic similarity despite different markers were used for their flow cytometric identification (Figure 2C). Taken together, the combined phenotypic and transcriptomic analysis presented here strongly argues for the need to further assess cellular identities within the myeloid cell compartment.

### Axl^+^Siglec6^+^ DCs phenotypically and transcriptionally overlap with human pre-DC

To clarify the relationships and cellular identities of the different DC subsets and their progenitors in maps 1 and 2, we mapped individual protein and transcript information (Figure S3) and the transcriptomic signatures of DC subsets and their progenitors derived from map 2 (pDC, cDC1, cDC2, pre-DC) onto our scRNA-seq myeloid-cell-space data set (Figure 3A). By overlaying index sorting information and the initial unbiased clustering data, we revealed that specific map 2 pDC, cDC1 or pre-DC signatures were enriched in dense discrete cell clusters within the UMAP topology of the myeloid-cell-space scRNA-seq data, whereas the cDC2 signature was more broadly enriched within both the clusters associated with cDC2 and monocytes (Figure 2A, 3A) suggesting a close relationship between these two cell types which is studied in further detail by Dutertre et al..

**Figure 3.**
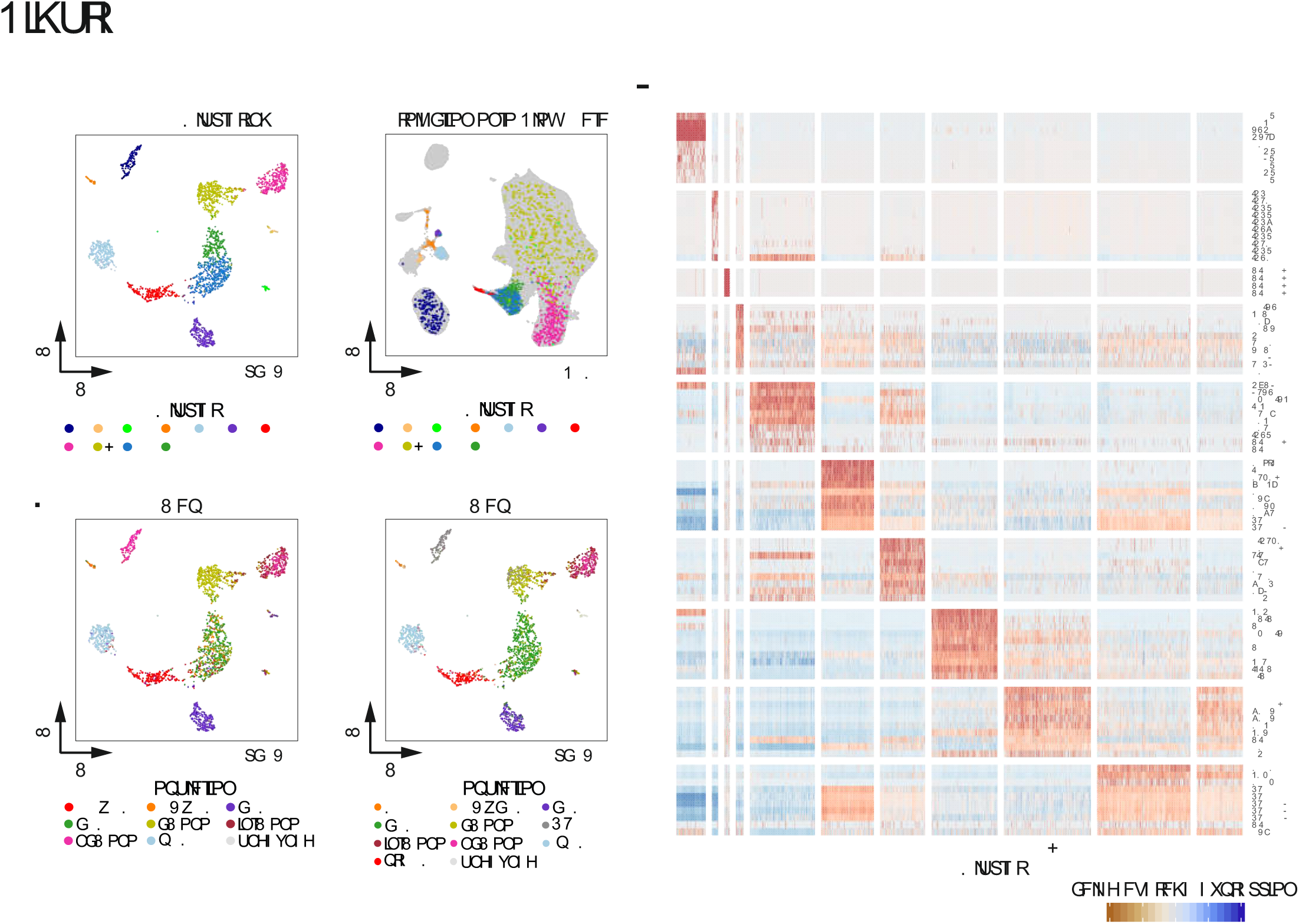
Harmonizing the DC space within the mononuclear myeloid cell compartment. (**A**) Overlay of signatures derived from map 2 DCs onto the new scRNA-seq-based UMAP topology consensus map. (**B**) UMAP topology based on map 1 single-cell transcriptomes of map 1 DC1-6 cells and overlay of signatures derived from map 2 DCs. (**C**) Overlay of signatures derived from map 1 DC1-6 cells onto the new scRNA-seq-based UMAP topology consensus map. UMAP topology based on flow cytometry data derived from ∼1.4 mio live CD45^+^Lin(CD3, CD19, CD20, CD56)^-^ cells (see Figure 1A) and separate overlays of cluster 26 defined by Phenograph (see Figure 1C), map 1 DC5, map 2 pre-DC, and scRNA-seq-based cluster seven representing transcriptomic progenitor DC signatures (see Figure 2A). **(E)** Enrichment of map 2 defined pDC, cDC1, cDC2 and pre-DC signatures in the map 1 DC1-6 subsets. **(F)** Heatmap of the average expression values of hallmark genes defined for map 1 DC1-6 subsets in both map 1 DC1-6 as well as map 2 DCs subsets.

To integrate the identified DC subsets in map 1 and map 2 with each other, we computed a UMAP topology from the original map 1 single-cell transcriptome data comprising the DC cell space and overlaid the signatures of the map 2 DC subsets (pDC, cDC1, cDC2, pre-DC) (Figure 3B). This analysis showed that within the original map 1 transcriptomic data, map 2 pDC signatures mapped to the same topological space as DC6, thus identifying DC6 as *bona fide* pDCs. Similarly map 2 cDC1 transcriptomic signatures were enriched within map 1 DC1, whereas map 2 cDC2 signatures enriched in map 1 DC2, DC3 and DC4. Furthermore, mapping map 2 pre-DC signatures revealed the highest enrichment of this signature in map 1 DC5 (AS-DC), indicating the highest level of similarity between map 1 DC5 and map 2 pre-DC.

To validate these correlations between the DC types defined in maps 1 and 2, we investigated the enrichment of map 1-defined DC1-6 signatures within our new scRNA-seq consensus data (Figure 3C). Visualizing the scaled signature enrichment scores for DC1 showed correspondence between maps 1 and 2 cDC1 locations and between map 1 DC2, DC3 and map 2 cDC2 locations, respectively. Similarly, map 1 DC6 and map 2 pDC localized to the same topological space within our new scRNA sequencing data. The highest enrichment of map 2 pre-DC signatures (Figure 3A) and map 1 DC5 signatures (Figure 3C) was seen in cluster seven of our new scRNA-seq consensus data (Figure 2A), again indicating substantial transcriptomic overlap between map 1 DC5 and map 2 pre-DC.

We then investigated the potential differences in cell-type identity between map 1 AS-DCs (DC5) and map 2 pre-DCs (Figure 2C). We separately projected cells identified as pre-DCs by unbiased clustering of the flow cytometric data (cluster 26 in Figure 1C), map 1 DC5, map 2 pre-DC gated cells and cluster seven from our new scRNA-seq consensus data, which displayed precursor gene expression patterns, onto the novel combined flow cytometric-based UMAP topology (Figure 3D). This approach showed that FACS cluster 26 represented the intersection of map 1 DC5, map 2 pre-DCs and scRNA-seq cluster 7 and best reflected these progenitor cells at the protein level in an unbiased fashion. Certain differences between map 1 AS-DCs and map 2 pre-DCs, however, became visible. Specifically, map 1 DC5 located only to a very discrete part of the topology and reached into a contaminating cDC2 space. FACS cluster 26 and map 2 pre-DCs occupied almost identical topological locations within the UMAP space, further illustrating the difficulties in discriminating pre-DCs and pDCs (Figure 3D, 1C-D, 2A). Interestingly, both map 1 AS-DCs and map 2 pre-DCs were best defined by FACS cluster 26, indicating that these cells represent the same cellular identity at both the surface marker and transcriptomic level. This finding was further supported when enriching transcriptomic signatures of map 2 pre-DCs across the spectrum of identified DC subtypes in map 1, resulting in a high enrichment of map 2-derived pre-DC signature genes within map 1 AS-DCs (Figure 3E). This enrichment was further reinforced by comparing hallmark genes within the cell populations defined in the legacy maps 1 and 2 (Figure 3F). In conclusion, these analyses demonstrate that map 1 DC5 and map 2 pre-DCs represent, to a large extent, the same pre-DC identities and therefore, might be best named according to already published guidelines (Guilliams et al., 2014; Schlitzer and Ginhoux, 2014) as pre-DCs.

### DN-DCs/DC4 resemble CD16^+^ non-classical monocytes

We were unable to locate the novel map 1 DC4 (DN-DC) subtype within a distinct cluster in our new scRNA-seq consensus data (Figure 2C). According to map 1 DC4 derived from a DN-DC subtype, being negative for the classical cDC subset markers CD1c, CD141 and CADM1 and pDC marker CD123 but positive for CD11c (Villani et al., 2017). To understand the role and placement of DC4 within the entire monocyte and DC space of both maps, we recapitulated the gating strategy originally used to delineate DC4 by map 1 (Figure 4A). Using the additional information from the newly included surface markers, such as CD16, we revealed that the large majority of DC4/DN-DCs (96.6%) were CD16^+^ mononuclear cells (Figure 4A). We subsequently mapped the CD16^-^ and CD16^+^ fraction of the DC4/DN-DC compartment of map 1 onto our integrated flow cytometry-derived and scRNA-seq-derived UMAP topologies (Figure 4B). Mapping onto the flow cytometry and scRNA-seq-derived UMAP topologies revealed that the CD16^+^ DC4/DN-DC compartment was associated with the location traditionally occupied by ncMono and the CD16^-^ fraction mapped into the topological region of the UMAP associated with pre-DCs and cDCs on both the phenotypic and transcriptomic level. To address the ambiguous DN-DC identity, we cross-referenced map 1 DN-DCs towards map 2 ncMono and the flow cytometry-based Phenograph cluster 15 (Figure 4C). Here we detected map 1 CD16^+^ DN-DCs almost exclusively within the map 2 ncMono cluster and primarily contained within Phenograph cluster 15 (Figure 1C) derived from the combined flow cytometry panel, expressing ncMono-associated surface markers.

**Figure 4.**
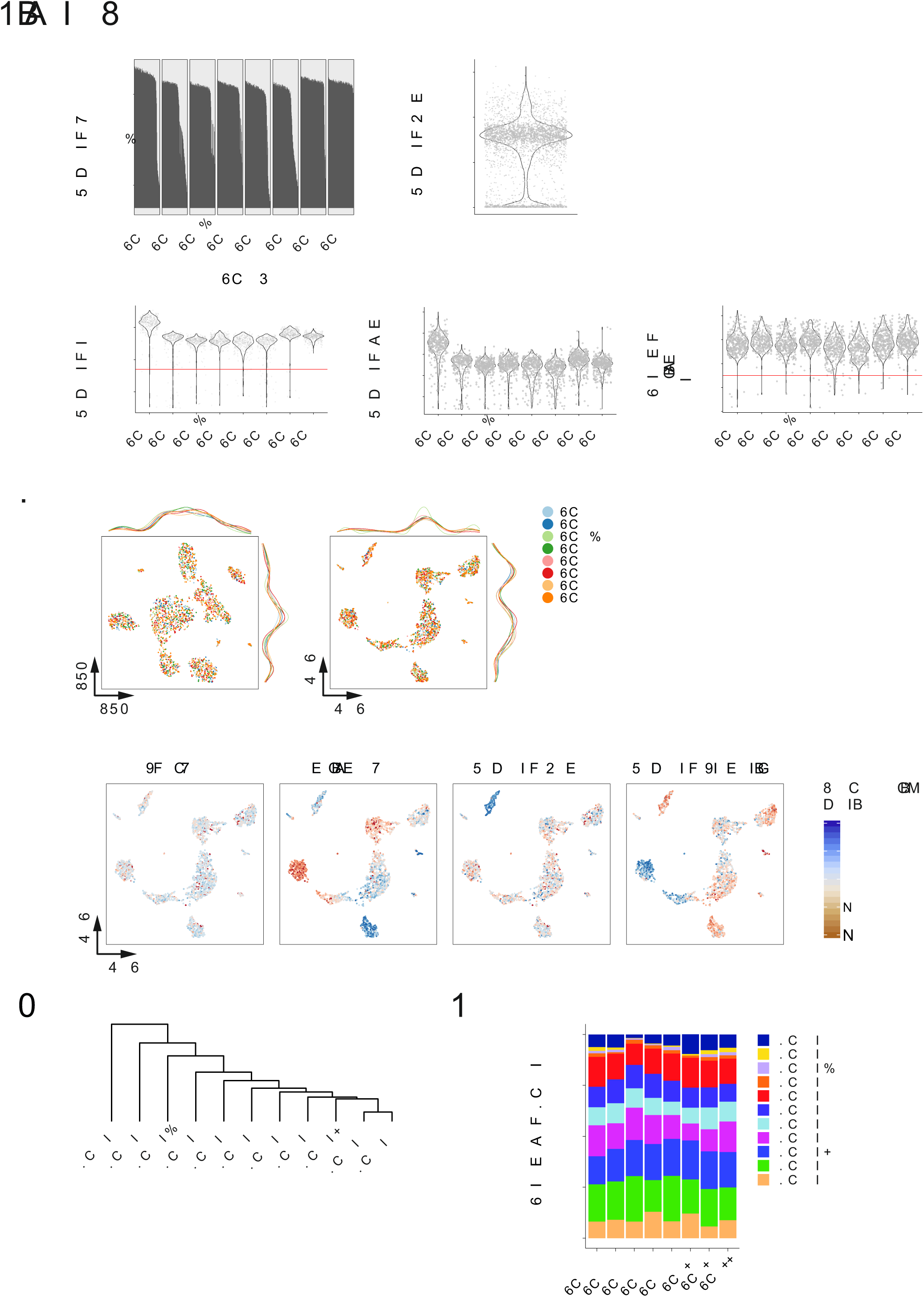
Integrating newly defined DN-DCs into the new consensus space of the myeloid cell compartment. (**A**) Recapitulation of the map 1 gating strategy to identify a putative DC4 subset within CD11c^+^ DN-DCs and visualization of CD16 expression. (**B**) CD16^+^ and CD16^-^ DN-DC mapping (as in A) onto the flow cytometry derived UMAP topology (left panel, see Figure 1B, 3A) and scRNA-seq data (right panel). (**C**) Relationship analysis of CD16^+^ or CD16^-^ DC-DN cells (see Figure 3A) and their corresponding annotation according to map 2, FACS Phenograph clustering and scRNA-seq data. (**D**) Pearson correlation matrix of all cell types defined in map 1. (**E**) Signature enrichment analysis of map 1 mono 2 signature in all other map 1-defined cell types. (**F**) Enrichment of the DC4 (DN-DC) signature visualized on the scRNA-seq data derived UMAP topology. (**G**) Gating strategy to define SLAN expression on the cell population defined as DN-DC based on CD16 expression. (**H**) Visualization of DN-DC cells in the complete CD45^+^Lin^-^HLA-DR^+^ UMAP topology (grey). (**I**) Mapping of the phenotypic information of cell populations onto the new UMAP topology.

To extrapolate these surface phenotypic findings to the transcriptome level and understand the transcriptomic identity of map 1 DN-DCs, we correlated all transcriptomes of map 1 DC subsets with the map 1 monocyte subsets (mono 1-4) (Figure 4D). Pearson correlation revealed the highest level of correlation between DC4 and the map 1 mono 2 subset, with intermediate correlation with the mono 1, 3 and 4 subsets, and poor correlation with any map 1-identified DC subset (Figure 4D). Furthermore, enrichment of a mono 2-specific gene signature across all map 1-identified mononuclear cell identities showed enrichment in all monocyte-associated cell entities and DC4, further supporting that DC4 might be ncMono (Figure 4E, S4A-C). Additionally, we used map 1-derived DC4 signature genes and mapped them onto our scRNA-seq consensus data of the blood myeloid cell space (Figure 4F). This analysis showed a strong enrichment of DC4 signature genes within the cluster identified by unbiased cluster detection as having ncMono identity.

To reconcile DC4 with the existing spectrum of monocyte and DC subsets, we examined DC4 expression of SLAN — a marker for inflammation-associated ncMono (Hansel et al., 2011) using a new marker panel (Figure 4G) and UMAP-based visualization (Figure 4H). DC4 showed the expected SLAN expression levels for ncMono. To validate this finding and to exclude that DC4 are another subset within peripheral blood mononuclear cells (PMBCs) that we might not have accounted for, we performed dimensionality reduction of the flow cytometric analysis in Figure 4G (Figure 4H) and mapped both CD16^-^ and CD16^+^ DN-DCs onto the UMAP topology (Figure 4I). Again, we found that DC4 associated with two different positions within this UMAP topology. Putting these two separate clusters within the DC4/DN-DC in the context of a conventional gating strategy of PBMC-derived mononuclear cells revealed a co-association between (i) classically defined ncMono and DC4/DN-DC that are CD16^+^ and constitute the already known ncMono fraction (Schakel et al., 1999), and (ii) a CD16^-^ pre-DC contamination associating with the areas within the UMAP defined as cDC2 and pre-DC by traditional investigator-informed gating (Figure 4I). Taken together, our new consensus map clarifies that map 1 DC4 is comprised of CD16^+^ ncMono and pre-DCs, rather than a phenotypically defined novel cell type within the human mononuclear myeloid cell compartment (Calzetti et al., 2018).

### Backmapping identifies mono 4 as bona fide CD56^dim^ NK cells

Next, we wanted to use our new consensus map to define the monocyte population structure. In particular, we aimed to consolidate the newly defined map 1 subtype structure with the four monocyte subtypes (mono 1-4) (Villani et al., 2017) in light of the traditional view of only three phenotypically different monocyte subsets based on CD14 and CD16 expression (Ziegler-Heitbrock et al., 2010). As a first step, we recapitulated the map 1 flow cytometry sorting strategy and overlaid the cellular contents of this gate onto our novel flow cytometry-derived UMAP topology (Figure 5A). Within the CD16^+^ compartment of map 1, two different monocyte populations (mono 2 and 4) were defined by phenotypical and transcriptional differences. Mapping the CD16^+^CD14^-^ compartment of map 1 onto the new UMAP topology indeed showed that it is composed of two transcriptionally different cellular entities, one mapping into the HLA-DR^-^ space of the flow cytometry-derived UMAP topology and one mapping to the location occupied by ncMono in an investigator-driven gating approach and named mono 2 in map 1 (Figure 1E, 2A, S1).

**Figure 5.**
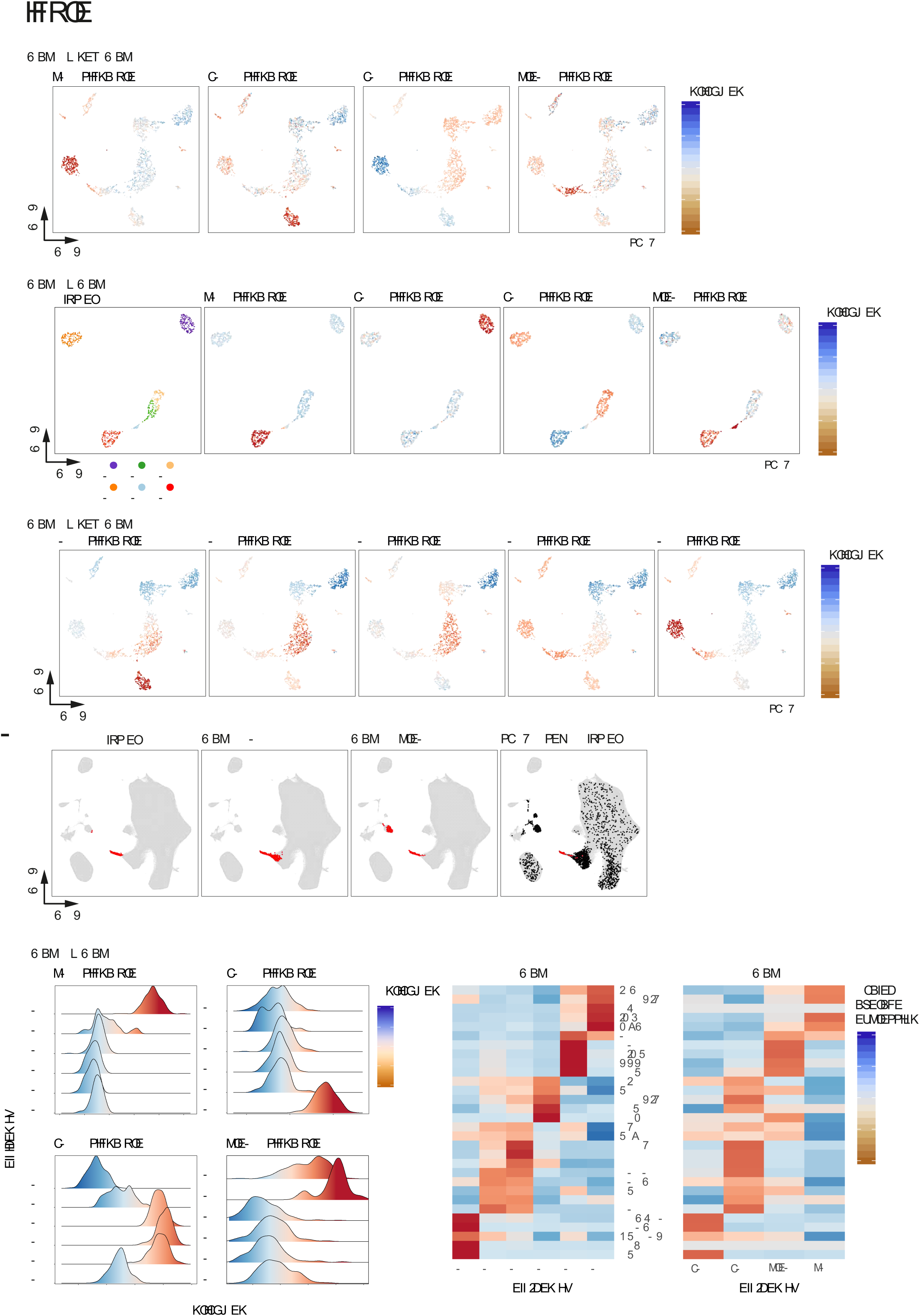
Relationship of the previously introduced fourth monocyte subset (mono 4) in context of the new consensus map of the myeloid cell compartment. (**A**) Mapping flow cytometric- and scRNA-seq-defined map 1 mono 2 and 4 onto the flow cytometry based UMAP topology (see Figure 1B, E, 2A). **(B)** Heatmap of the protein expression pattern for mono 4 signatures genes derived from mass-spectrometric data of FACS-sorted population as described by Rieckmann et al. (Rieckmann et al., 2017). (**C**) Enrichment of a NK cell signature in map 1 mono 1-4 subsets. (**D**) Heatmap of NK cell hallmark genes within the map 1 defined cell subsets. (**E**) Backmapping by overlaying map 1 mono 1-4 cells onto the 33k-PBMC scRNA-seq dataset. Only monocytes and NK cells are shown. (**F**) Visualization of the percentage of cells that are aligned with either monocytes or NK cells derived from the unrelated 33k-PBMC scRNA-seq dataset. (**G**) UMAP topology of scRNA-seq data derived from the map1 DC and mono subsets (left panel) and overlay of the NK cell signature onto this UMAP topology. (**H**) Top panels: classical gating strategy and stepwise cleanup of CD45^+^ cells based on lineage (CD3/CD19/CD20) marker expression, then based on CD56 expression followed by HLA-DR expression, left to right. Middle panels: UMAP topology derived from the respective cell populations marked within the corresponding top panels. Mono/DC by HLA-DR expression, green; NK by CD7 expression, violet; and granulocytes by CCR3 or CD66b expression, orange. Bottom panels: Effect of cleanup as shown in top panels on the CD16^+^ CD14^-^ cell population.

To understand the identity of the cells mapping to the HLA-DR^-^ cell space, we utilized the transcriptomic marker genes derived from map 1 mono 4, as mono 2 mapped to the ncMono space. We then interrogated a publicly available database of population-based proteome fingerprints (Rieckmann et al., 2017) from a variety of blood-borne immune cells for the mono 4 signature (Figure 5B). Here, we found high expression of mono 4-related proteins in NK cell subsets, including a CD56^dim/neg^ subset (NK^dim^). To validate these findings, we generated a transcriptomic NK-cell signature based on previous knowledge (Costanzo et al., 2018; Liberzon et al., 2011; Rieckmann et al., 2017; Subramanian et al., 2005) and calculated the signature enrichment scores across all monocyte subsets defined in map 1 (Figure 5C-D). This calculation revealed that the mono 4 subset was significantly enriched in NK-cell-specific transcripts. Subsequently, we integrated the original monocyte map 1 single-cell transcriptome data (mono 1-4) into an external dataset of 33,148 PBMCs (short: 33k-PBMC dataset, https://support.10xgenomics.com/single-cell-gene-expression/datasets/1.1.0/pbmc33k) and performed dimensionality reduction of the corresponding monocyte and NK-cell-related cellular spaces using UMAP (Figure 5E-F). We termed this approach ‘backmapping’, where we utilized an unrelated single-cell data set derived from the same tissue origin. In a next step, we annotated the integrated map 1 specific monocyte subsets within the combined UMAP topology according to the terminology used in map 1, to understand where these cells would associate in the context of an unbiased assessment of the complete mononuclear PBMC fraction. This analysis showed that the mono 1-3 subsets mapped to the topological UMAP space initially assigned to monocytes, whereas mono 4 mapped to the topological UMAP space of NK cells, further supporting the hypothesis that mono 4 are NK cells. Overlaying of the NK cell signature onto the original map 1 also revealed strong enrichment in the mono 4 cluster (Figure 5G).

We then modified our combined flow cytometry panel to specifically verify NK cell contamination within the map 1-defined flow cytometry CD16^+^ monocyte cell space (Figure 5H, S5A). Specifically, we removed CD56 from the lineage to track the expression of this NK-cell marker separately and added the granulocyte marker CD66b, the lymphoid marker CD7 and the NK cell markers NKp46, CD160 and CD107 (Figure S5A). We then examined CD16 and CD56 expression within the CD14^-^ compartment of the PBMC CD45^+^Lin^-^ fraction and generated a reference UMAP topology of the CD45^+^Lin^-^ cell space (Figure 5H, S5A-B). This analysis identified seven cell populations based on CD16 and CD56 expression levels (Figure S5B-C). Two populations (turquoise and pink) displayed high CD16 and SSC and no (light blue) to mid (pink) CD56 expression, with granulocytic forward and sideward scatter characteristics identifying them as granulocyte contaminants (Figure S5C).

Next, we focused our analysis on the CD16^+^ compartment of this cell space (green and purple gates). Two populations were identified as CD16^+^, in which one CD16^+^CD56^+^ population (purple) matched the surface phenotype of classical CD56^+^ NK-cells that are normally dismissed by including CD56 in the lineage panel of map 1. To determine the identity and heterogeneity of the remaining CD16^+^CD56^-^ cell compartment (green, orange, yellow, grey gates, Figure S5B), we mapped this compartment back to a UMAP topology of either the Lin^-^CD16^+^, Lin^-^CD56^-^CD16^+^ or the Lin^-^CD56^-^CD16^+^HLA-DR^+^ cell space, to represent a stepwise cleanup of non-monocytic CD16^+^ cells (Figure 5H). This analysis showed that if the totality of the Lin^-^CD16^+^ compartment is mapped back onto the Lin^-^ UMAP topology (Figure 5H, pink overlay, most left panel), NK cells (CD56^+^), monocytes (CD56^-^CD16^+/-^) and granulocyte fractions (CD16^high^) are included in this cellular compartment. When excluding CD56 in the UMAP topology, classical CD56^+^ NK cells are excluded; however, within the CD16^+^ gate a CD56^-^ population became apparent that mapped to the UMAP space previously associated with classical NK cells (Figure 5H, pink overlay, mid panel). Another CD16+CD56^-^ population mapped to the topological UMAP space occupied by ncMono, as defined by their high expression of HLA-DR and CD11c and no expression of classical NK-cell markers, including CD56, CD7, CD160 and Nkp46 (Figure S5C). We next excluded HLA-DR^-^ cells and mapped the remaining CD16^+^ cells onto the UMAP topology. This step revealed that including a positive HLA-DR threshold successfully removed mono 4 / NK cells (pink overlay, Figure 5H, right panel). This subclass of NK cells is not easily distinguishable from monocytes, as also evidenced by their very similar morphology (Figure S5D). Taken together, we identified the map 1 mono 4 subset as HLA-DR^-^CD16^+^CD56^-^ NK-cells, intruding into the CD16^+^CD14^-^Lin^-^ CD45^+^ map 1 monocyte sorting gate that was performed without HLA-DR gating stratification according to map 1. Collectively, within our new consensus map, we define the borders between the myeloid compartment and the NK cell compartment and re-stablish a structure of three monocyte subsets in peripheral blood.

### PBMC derived monocyte subsets form a transcriptional continuum during homeostasis

*De novo* clustering (Figure 2A, 6A) did not reveal intMono as a transcriptionally distinct cluster; rather, they were identified as forming part of clusters eight and nine (Figure 6A, 6B, S6A, S6B). Pseudo-time analysis of the scRNA-seq data, however, placed intMono in between cMono and ncMono (Figure 6C). The visualization of genes changing over the pseudo-time depicts a gradual decrease in expression of cMono marker genes (CD14, etc) and an increase of ncMono marker gene expression (CD) along the trajectory (Figure 6D). This was further corroborated by plotting CD14 and CD16 mRNA expression of single cells within the three monocyte subsets (Figure 6E). Therefore, these analyses clearly corroborate an existing transcriptional continuum of monocytes within human PBMC and reveal the transcriptional identity of intMono during homeostasis.

**Figure 6.**
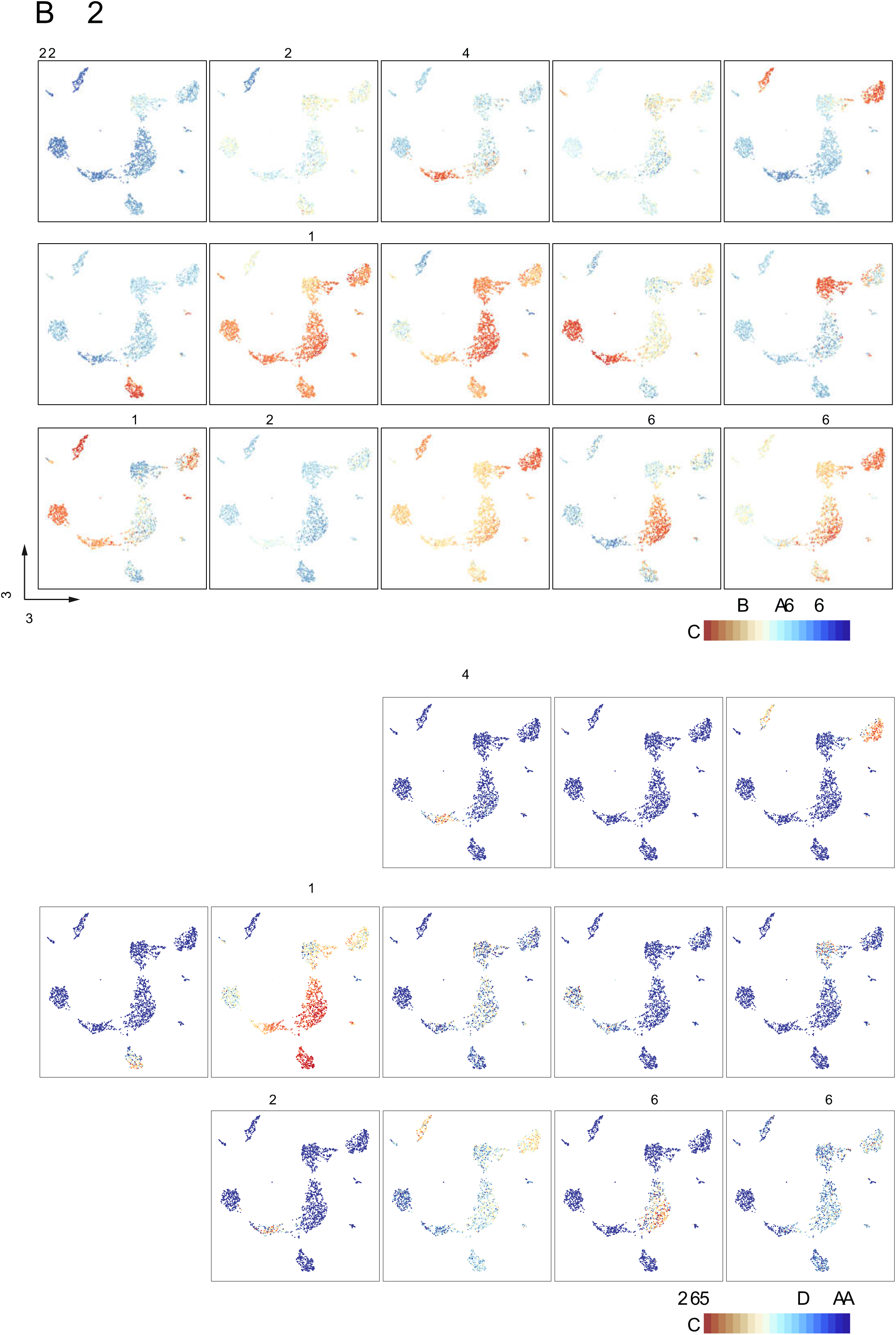
Focused analysis of the monocyte compartment. **(A)** Overlay of the the cluster 8 and 9 defined by de novo clustering of the scRNA-Seq data onto the scRNA-UMAP topology of the new consensus map. **(B)** Bar chart showing the original FACS annotation of cells derived from cluster 8 or 9 following the sorting scheme of map 1 or map 2, respectively. **(C)** Trajectory analysis of the monocyte subset containing cells from cluster 8 and 9. Monocle-based UMAP dimensionality reduction overlaid with cell estimated pseudo-time (left panel) and the FACS annotations derived from map 1 (center panel) or map 2 (right panel). **(D)** Transcriptional changes of genes that are considered as differentially expressed along the inferred trajectory. Heatmap shows scaled gene expression changing over the pseudo-time (x-axis, early to late). Selected marker genes of cMono and ncMono are highlighted. **(E)** Expression of CD14 and CD16 in relation to the estimated pseudo-time of cells. Cells are colored by their FACS annotation from map 1 or map 2.

### Backmapping integrates legacy datasets and enhances cell type resolution creating novel consensus maps

To evaluate our findings and put our new consensus map into the framework of data-driven maps of the complete human PMBC compartment, we combined our new scRNA-seq data with three independent PBMC datasets and performed backmapping and cell type prediction (Figure 7, S7). This approach permitted a detailed annotation of previously undefined cellular identities within the external PBMC-derived datasets. *De novo* analysis of a combined data set (Figure S7A) consisting of a publicly available dataset of approximately 33.000 cells (33k-PBMC) and our new scRNA-seq dataset revealed NK, CD8^+^, CD4^+^ and B cells alongside megakaryocytes, CD16^+^ and CD14^+^ monocytes, cDC1, DC2, DC3, pDCs and CD34^+^ progenitor cells (Figure 7A). When using the new scRNA-seq-based consensus map as a reference to predict cell annotations in the data-driven PBMC dataset by a nearest neighbor classifier (Kiselev et al., 2018), we obtained similar but not identical results (Figure 7B, left panel). We therefore applied the backmapping approach by projecting the cluster identities of our new scRNA-seq dataset onto the UMAP dimensionality reduction of the combined dataset. Backmapping revealed commonalities between all cell types of the myeloid cell compartment including previously unidentified pre-DCs, DC2 and DC3 clusters within the 33-K PBMC dataset (Figure 7B) as well as NK cells, CD34^+^ and plasma cells derived from both datasets (Figure S7B). We next applied the above outlined approach including backmapping to a larger data set provided by the Human Cell Atlas (HCA) with approximately

**Figure 7.**
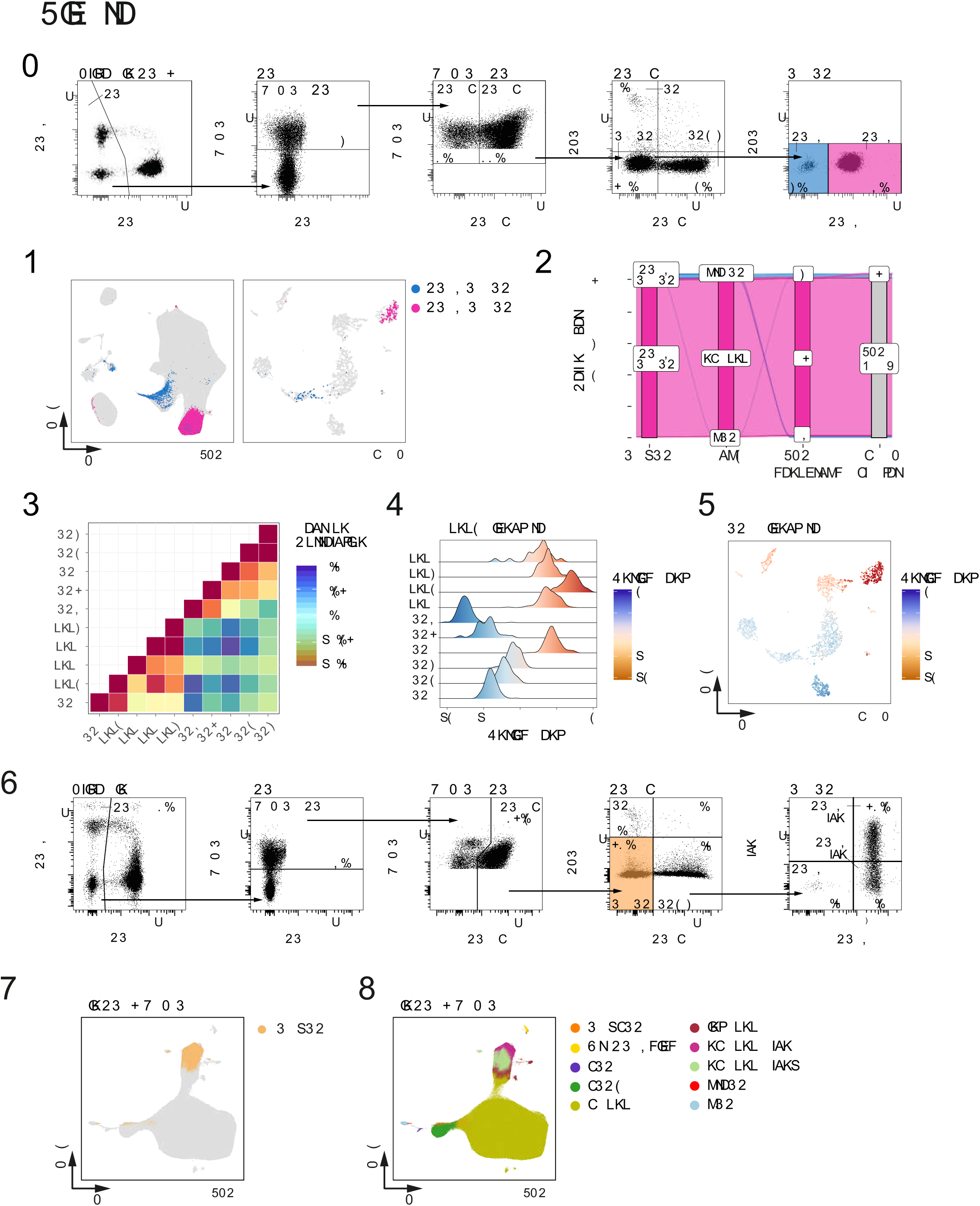
Backmapping strategy combining the new scRNA-seq data and different PBMC datasets. **(A)** Annotation of cell types within the combined dataset (33-K PBMC and new scRNA-seq dataset). **(B)** Graphs in the left panel predict cell labels from the 33-K PBMC dataset by using the transcriptome information from the new scRNA-seq dataset. Graphs in the right panel show the visualization of the cells from the new scRNA-seq dataset after integration with the 33-k PBMC dataset. **(C)** UMAP dimensionality reduction of around 260.000 human cord blood cells and cell annotation based on markers obtained from the unrelated 33k-PBMC dataset (Figure 7A-B). **(D)** Reduction of the HCA dataset to cells, which were found within clusters associated with monocytes or dendritic cells. **(E)** Graphs within the left panel show the prediction scores calculated for the respective cell types of the new scRNA-seq data. Graphs in the right panel show the visualization of the cells from the new scRNA-seq data after “anchoring” together with the HCA dataset.

255.000 cells (Figure 7C, 7D, 7E). In a first step, we categorized all major subtypes (Figure 7C) and predicted cells associated with the mono 4 population of map 1 and found that all cells fell within the NK cell cluster (Figure S7C, S7D), further supporting that cells of the mono 4-subset are *bona fide* NK cells. Next, we reduced the datasets to clusters that were part of the myeloid cell compartment (Figure 7D, S7C) and again performed the backmapping and prediction approaches. Because of the solely cluster-driven reduction of the dataset, we observed some lymphoid cells in the reduced dataset, which might derive from misclustered cells (Figure 7D). Nevertheless, this approach allowed us to identify smaller myeloid cell populations within the larger dataset (Figure 7E). Finally, we used a third PBMC dataset based on a targeted scRNA-seq approach, which also allowed us to better define subsets in the unbiased dataset (Figure S7E, S7F, S7G) making this overall approach independent of single-cell technology and dataset size.

Collectively, we demonstrate the value of an iterative, rule-based, data-informed approach based on previously existing maps to integrate additional information layers into new consensus maps. Backmapping to whole data-driven tissue maps and providing a connection to previous knowledge are important steps to derive the next iterations of maps that finally serve as the entry point for further iterations.

## Discussion

Consensus maps are an important instrument within an iterative process of producing cellular maps of all organs and tissues in different species, including humans. As within other scientific disciplines, such as geography or astronomy, the maps generated in the life sciences require much iteration to allow for the integration of new content. By combining single-cell transcriptomics with index sorting, and multi-color flow cytometry and applying simple but very effective computational strategies, such as ‘backmapping’ to cellular maps generated in a purely data-driven fashion, we have generated a new consensus map of the myeloid cell compartment including monocytes, DCs and their precursors (Figure 7, S7). Because we propose to include prior knowledge in the respective scientific field into the algorithm for generating such consensus maps, we define the overall strategy as being ‘data-informed’, combining prior knowledge and data-driven technologies including single-cell omics.

The two previous maps based on single-cell RNA-seq used in our approach as well as a phenotypic analysis of the human blood and tissue myeloid cells were developed to improve our understanding of myeloid cell heterogeneity (Alcantara-Hernandez et al., 2017; See et al., 2017; Villani et al., 2017). Yet there were shortcomings to these maps, which we have overcome in our new consensus map. First, map 2 only identified one cDC2 subset, whereas map 1 and our new consensus map defined two subsets. Furthermore, we established a proximity between DC3 and cMono, which has been further dissected by Dutertre *et al.* (Dutertre et al., 2019), thus already providing the next iteration of this particular subspace in the myeloid cell map of human peripheral blood. Second, map 1 identified a novel monocyte subset named mono 4. Using backmapping, we reveal that mono 4 are CD56^dim^ NK cells and are not related to monocytes, supporting the current definition of three major monocyte subsets (classical CD14^+^, non-classical CD16^+^, and intermediate double positive monocytes), consisting of two transcriptionally distinct entities and a continuum of intermediate, double-positive monocytes between them (Figure 6). This finding is further supported by the changes in expression of the CD14 and CD16 cell-surface markers and results derived from genetic mouse models showing that Ly6c^hi^ monocytes (murine equivalents of classical monocytes) can transition into Ly6c^low^ monocytes (murine equivalents of non-classical monocytes) with only a few cells detectable in the transitory state (Mildner et al., 2017). Third, we could clearly define AS-DCs (DC5) from map 1 as pre-DCs within the consensus map, consistent with their functional definition in map 2 (See et al., 2017). Together with the complete overlap of the three differentiated DC populations between the original maps, these results reassure the validity of single-cell transcriptomic analyses.

We define backmapping as an integral component of the strategy to define novel consensus maps. Here, we use cellular maps derived from tissues – in this case peripheral blood - without prior experimental enrichment of certain cell types. This relatively simple computational approach allows to unequivocally overlay cell subsets from different maps onto a common cell space. As exemplified here for the monocyte / NK cell space, we postulate that potential conflicts for new cell types in other organs can be resolved in a similar fashion.

Collectively, we report on a new consensus map of the myeloid cell compartment in human blood, which was built on two previously introduced maps (See et al., 2017; Villani et al., 2017). The myeloid cell compartment is of particular interest due to its intrinsic heterogeneity, its involvement in many if not all major tissues and organs, and its prime involvement in almost any major disease (Bassler et al., 2019). It is therefore of utmost importance to establish a precise baseline during homeostasis as provided here by our new consensus map, to allow for a better understanding of any deviations of the myeloid cell compartment during stress, pathophysiological conditions and diseases. Furthermore, the dynamic processes of myelopoiesis during homeostasis but even more so during inflammatory conditions requires precise mapping of cellular identities as a prerequisite to identify targets for precise therapeutic intervention (Dick et al., 2019; Schultze, 2019; Schultze et al., 2019). Furthermore, the necessity to continuously iterate the process of improving the consensus maps is nicely illustrated by the accompanying manuscript by Dutertre *et al.* (Dutertre et al., 2019), further defining the cellular relationship of DC2/3 and monocytes. As many institutions world-wide continue to generate cellular maps, consensus maps will become an increasingly important instrument to reconcile and integrate information. Our approach provides a guide to integrate and value legacy datasets together with newly generated single-cell omics data and build new iterations of consensus maps applicable to any other tissue. These maps can also be adapted to include further technological advancements. With the continuation of technical advances, we anticipate that consensus map building will become a major task within our efforts to create complete cellular atlases for the major species.

## Supporting information

Antibody panel

## Acknowledgments

*Non-author contributions:* We thank Jessica Tamanini for critical review and editing of the manuscript.

## Author contributions

Conceptualization, P.G, B.C., A.S. and J.L.S.; Methodology, P.G, B.C., A.S. and J.L.S.; Software, P.G, B.C, K.B., M.B. and K.H., Investigation, P.G, B.C, K.B., K.H., C.A.D. and V.B., Resources, A.S. and J.L.S., Writing – Original Draft, P.G, B.C., A.S. and J.L.S.; Writing – Review & Editing, P.G, B.C, K.B., E.N., M.C., F.G., A.S. and J.L.S.; Visualization, P.G, B.C. and K.B.; Supervision, A.S. and J.L.S.; Project Administration, A.S. and J.L.S.; Funding Acquisition, E.N., M.C., F.G., A.S. and J.L.S..

## Data and materials availability

Processed and raw scRNA-seq datasets are available through the Gene Expression Omnibus (GSE126422). In addition, we provide an interactive web tool to visualize the single-cell RNA-Seq data together with the flow cytometry data at https://paguen.shinyapps.io/DC_MONO/.

## Supplementary Figure Legends

**Figure S1.**
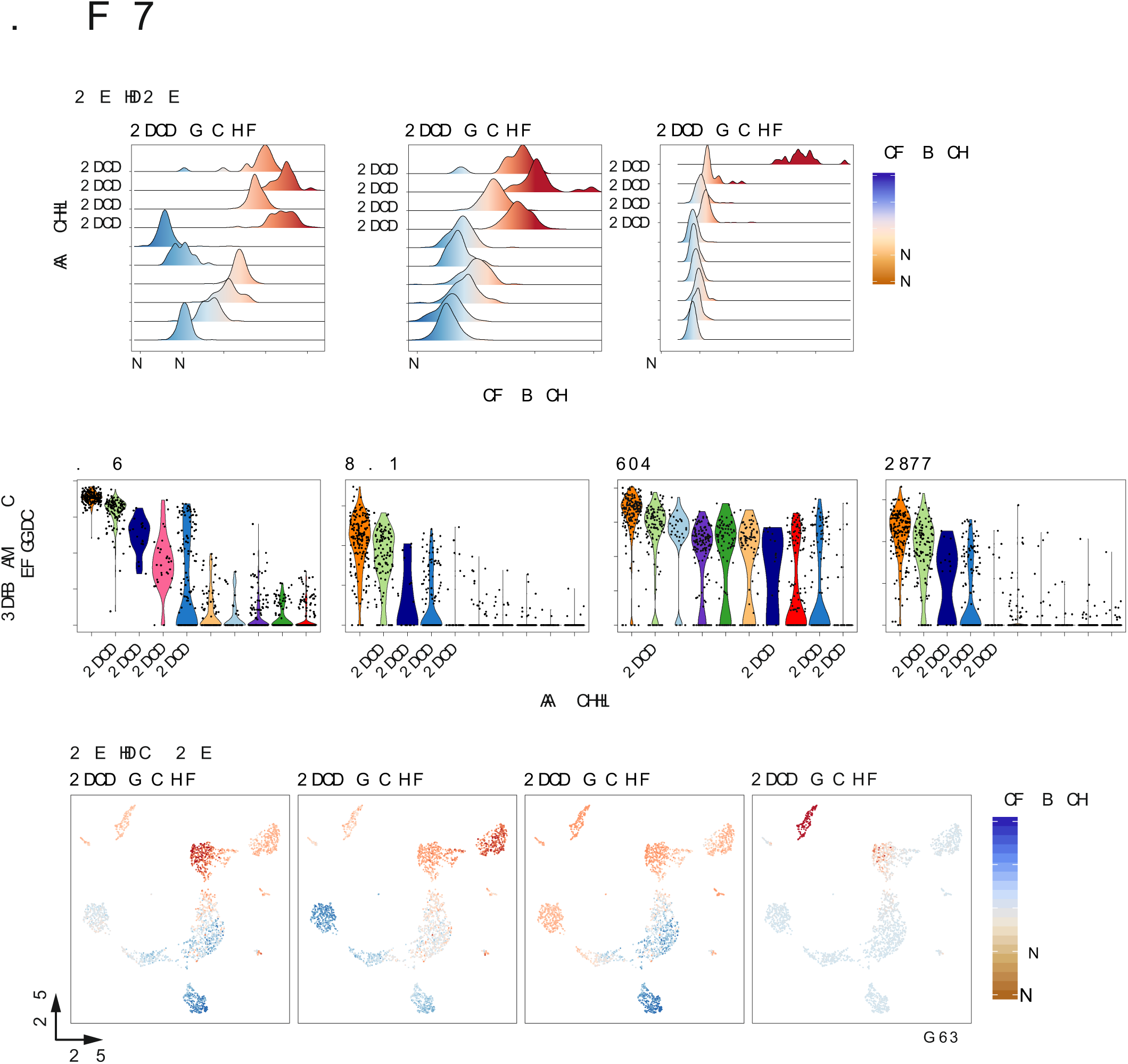
Classical flow cytometry gating strategies applied for the generation of the legacy maps 1 and 2. **(A)** Common part of the flow cytometry gating strategy applied for the generation of the legacy maps 1 and 2. **(B)** Map 1-specific part of the flow cytometry gating strategy with the resulting cell subsets (colored boxes). **(C)** Map 2-specific part of the flow cytometry gating strategy with the resulting cell subsets (colored boxes). Some cell types shown here are based on a priori definitions (e.g. monocytes) and were not part of the transcriptionally defined cells in the legacy map 2.

**Figure S2.**
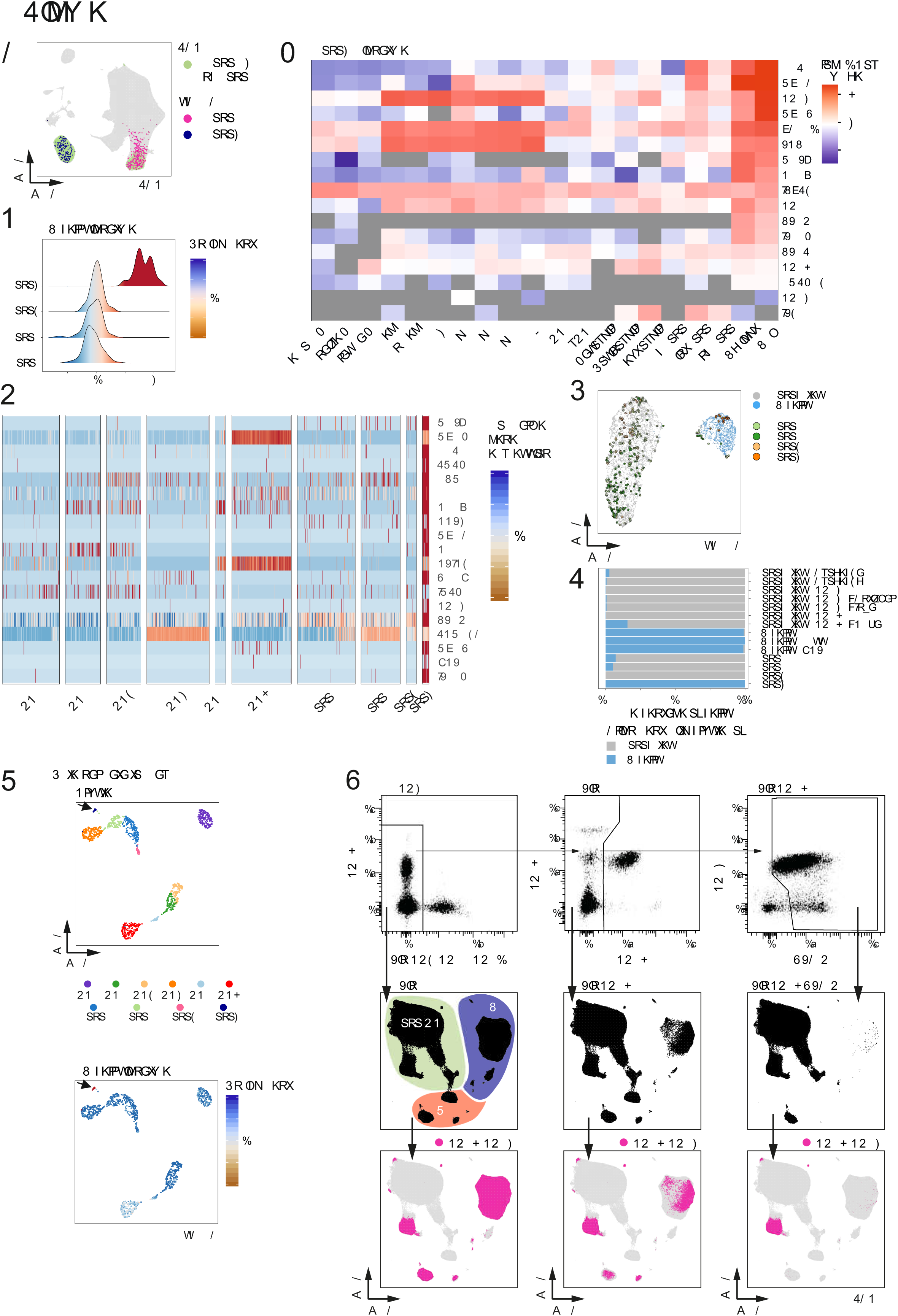
Quality control criteria for the scRNA-seq data established for the new consensus map of the myeloid cell compartment. **(A)** Visualization of the number of reads for all 8 384-well plates analyzed within this project. Violin plot of the number of genes observed to be present within all cells measured. **(B)** Visualization of the number of reads (left panel), the number of genes (middle panel), and the percent of aligned reads (right panel) as a violin plot for each of the 8 384-well plates individually. **(C)** Comparison and visualization of cell distribution across all identified clusters in relationship to the 8 384-well plates utilized within this experiment. **(D)** Mapping of single-cell information concerning the total number of reads, unaligned reads, number of genes, and number of transcripts onto the UMAP topology of the final consensus map based on scRNA-seq data. **(E)** Cluster relationship analysis **(F)** Distribution of cells within each of the identified cluster in relation to the 8 384-well plates used in this study.

**Figure S3.**
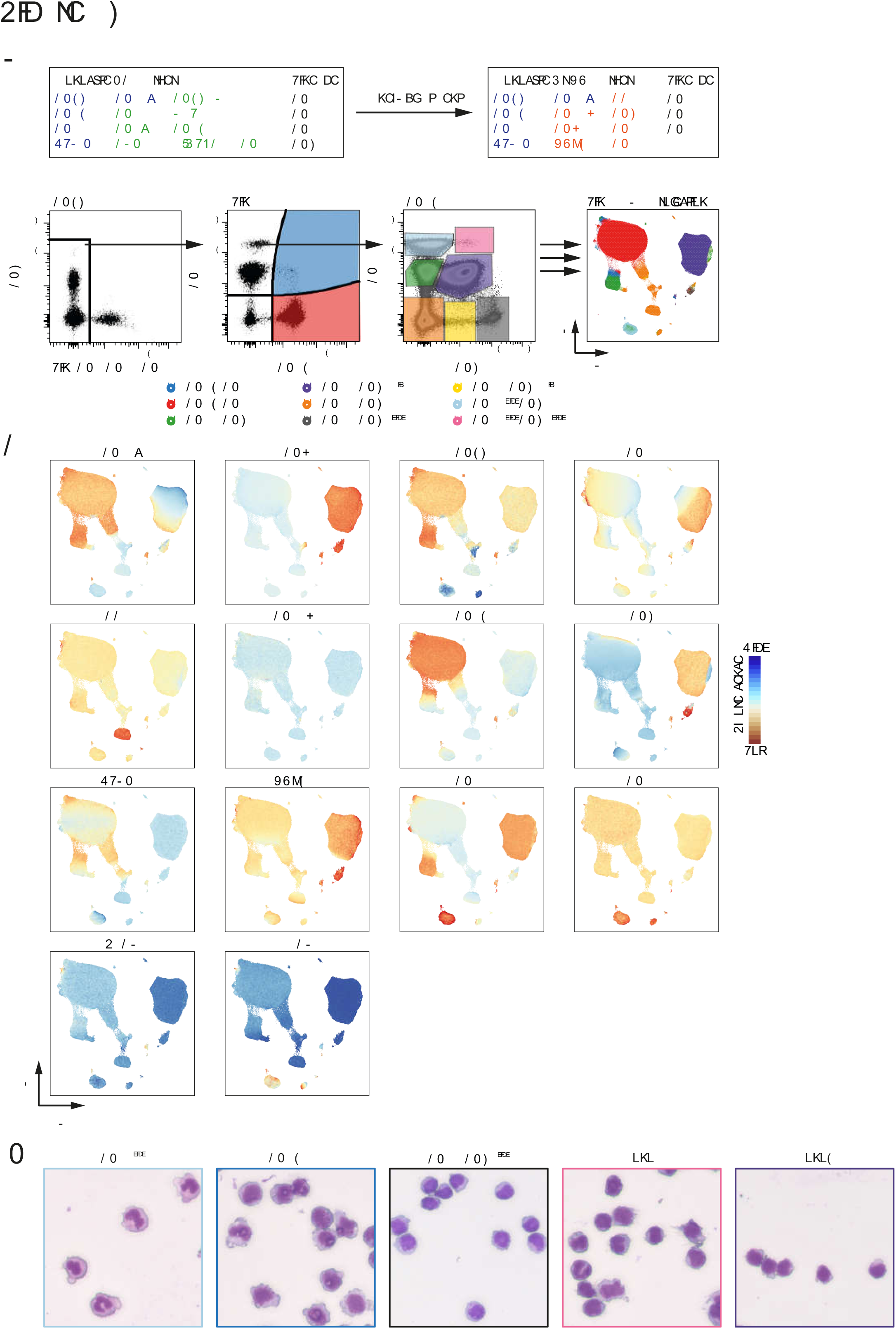
Overlay of phenotypic and transcriptomic data onto the new consensus map of the myeloid cell compartment. **(A)** Visualization of cell surface markers detected by index sorting on the 2,509 cells which were used to define the UMAP topology of the index-sorted cells based on the single cell transcriptome data. **(B)** Visualization of gene-level expression of the respective cell surface markers within the 2.509 cells which were used to define the UMAP topology of the index-sorted cells based on the single cell transcriptome data.

**Figure S4.**
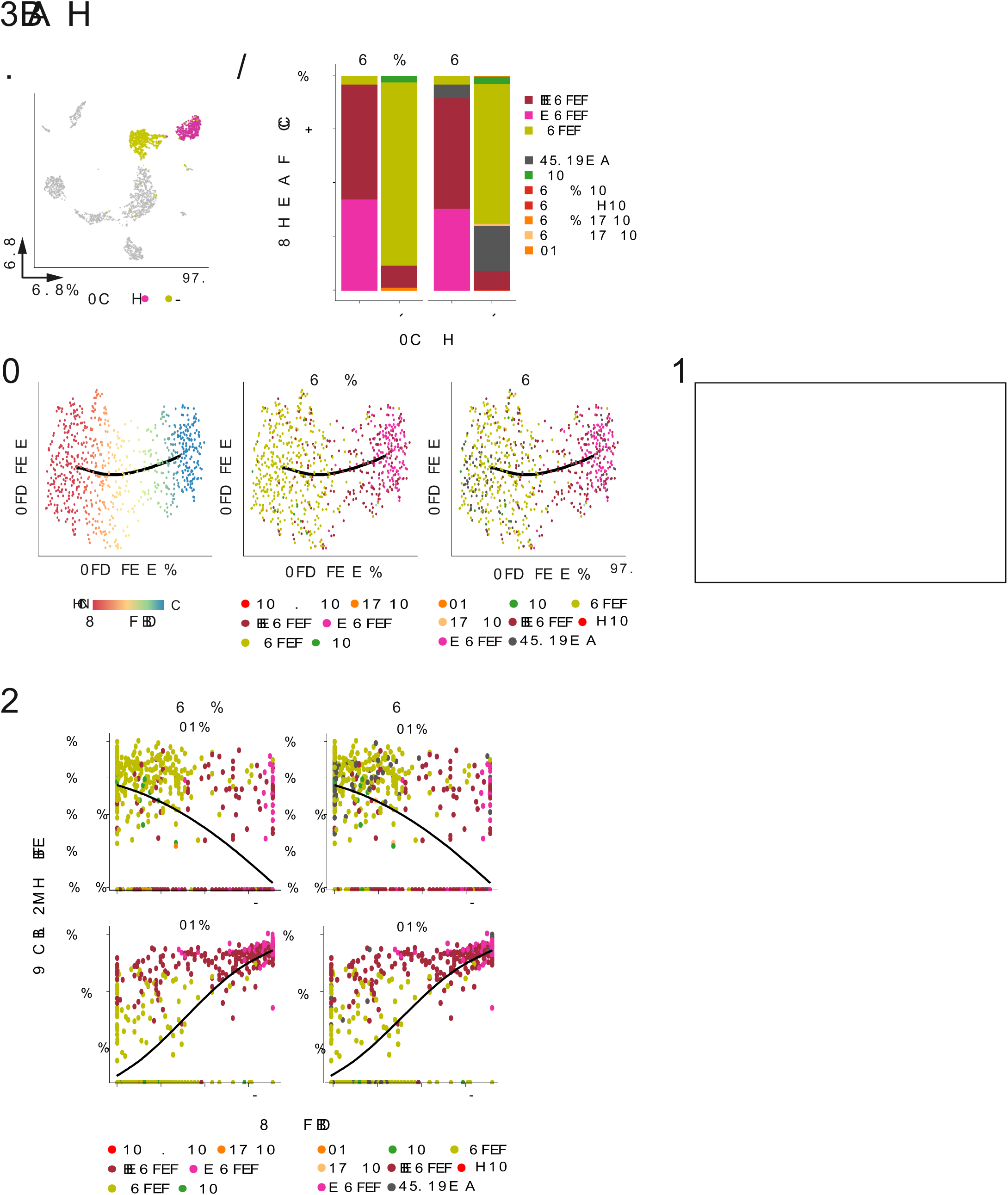
Defining the relationship of map 1 DN-DC within the new consensus map of the myeloid cell compartment. **(A)** Enrichment of map 1 mono 1,3,4 signatures in map 1 mono1-4 and DC1-6 subsets. **(B**) Violin plots for normalized gene expression of FCGR3A, TCF7L2, RHOC, MTSS1 in map 1 mono1-4 and DC1-6 subsets. **(C)** Overlay and visualization of map 1 mono 1-4 subset gene signature enrichment on the UMAP topology based on scRNA-seq of 2.509 index-sorted cells (see Figure 2A).

**Figure S5.**
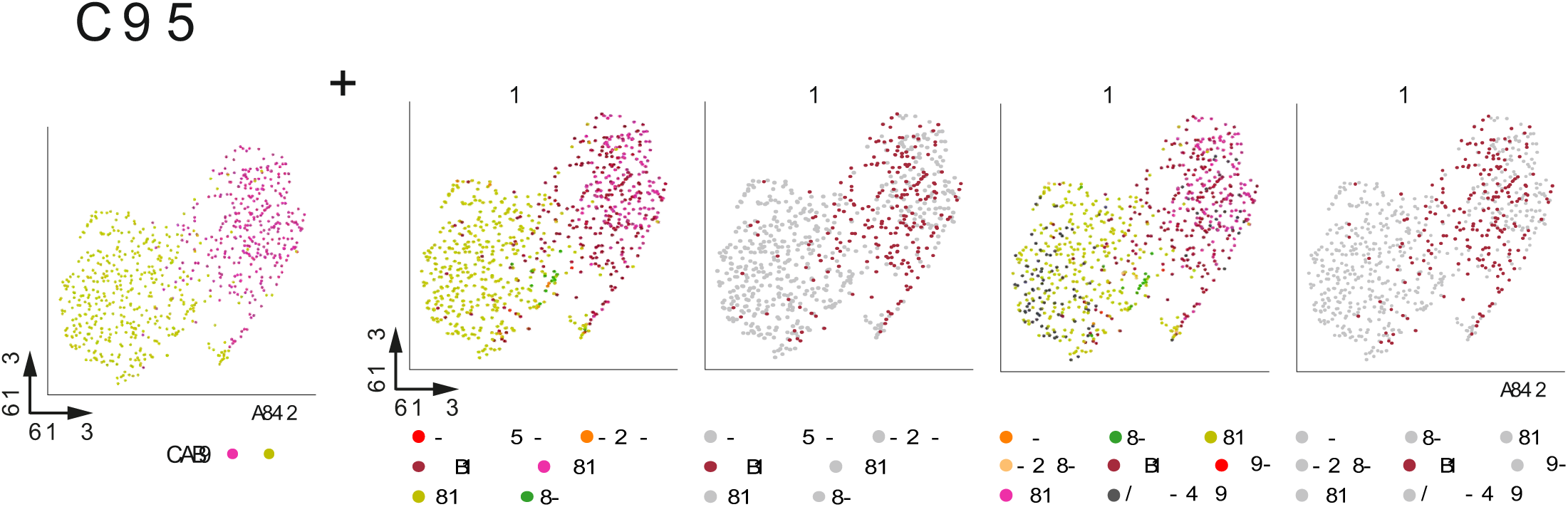
The relationship between the myeloid and the NK cell compartment in human peripheral blood. **(A)** Schematic representation of the development of a new focus strategy (panel adjustment) to define the relationship between the myeloid and the NK cell compartment in human peripheral blood. **(B)** Classical gating strategy to determine those cell populations that need to be placed either into the myeloid or the NK cell compartment followed by the development and visualization of the UMAP topology of both cellular compartments. **(C)** Color-coded visualization of markers used to define the mononuclear myeloid and NK cell compartments on the flow cytometry data-based UMAP topology. **(D)** Cytospins of cells sorted according to the gating strategy depicted in Figure S5B.

**Figure S6.**
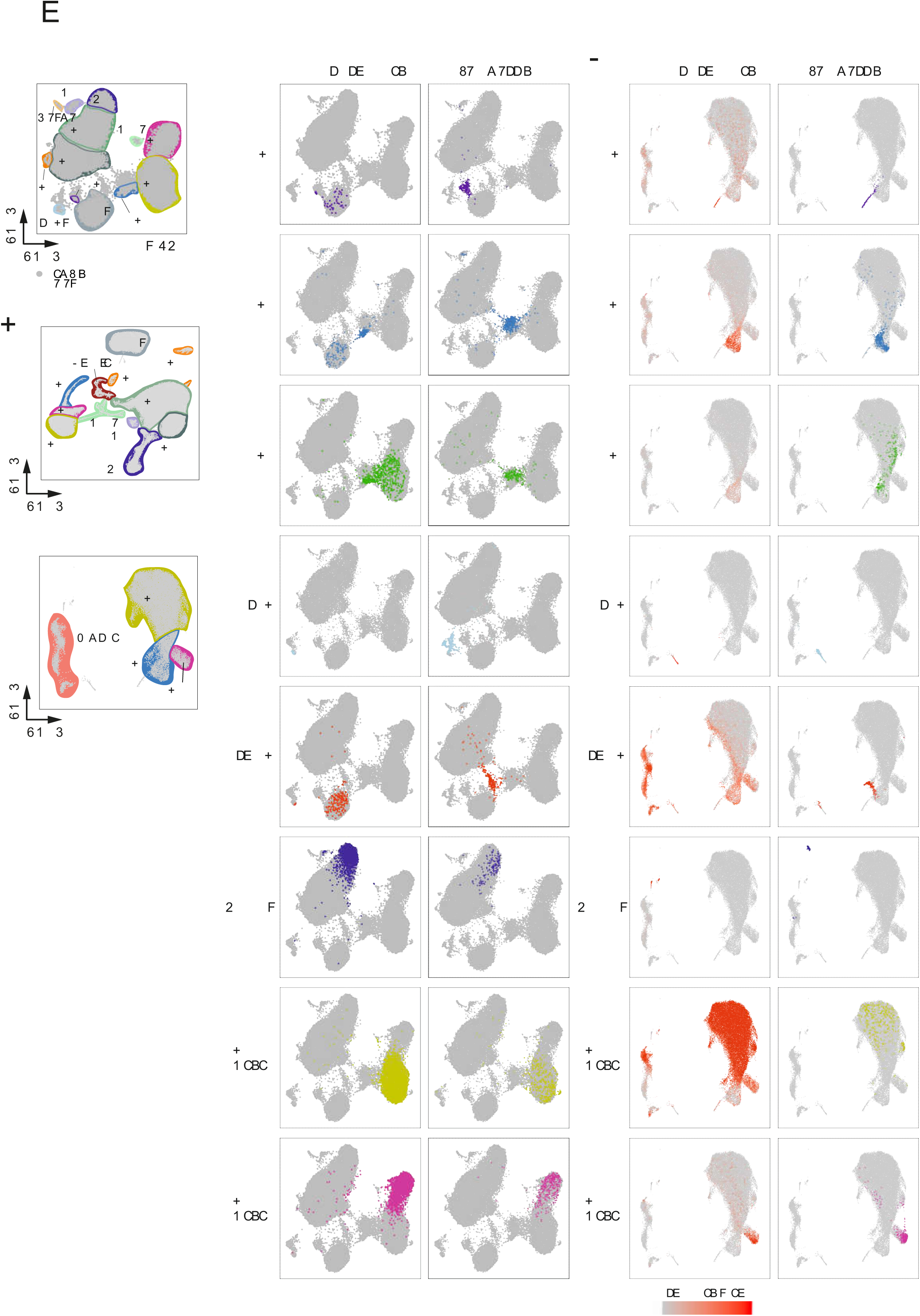
Focused analysis of the monocyte compartment. **(A)** UMAP dimensionality reduction calculated on the focused subset of cluster 8 and 9 containing the monocyte populations. Overlaid are the cell annotations from the clustering and **(B)** the FACS annotation derived from Map 1 (upper panel) and Map 2 (lower panel).

**Figure S7.**
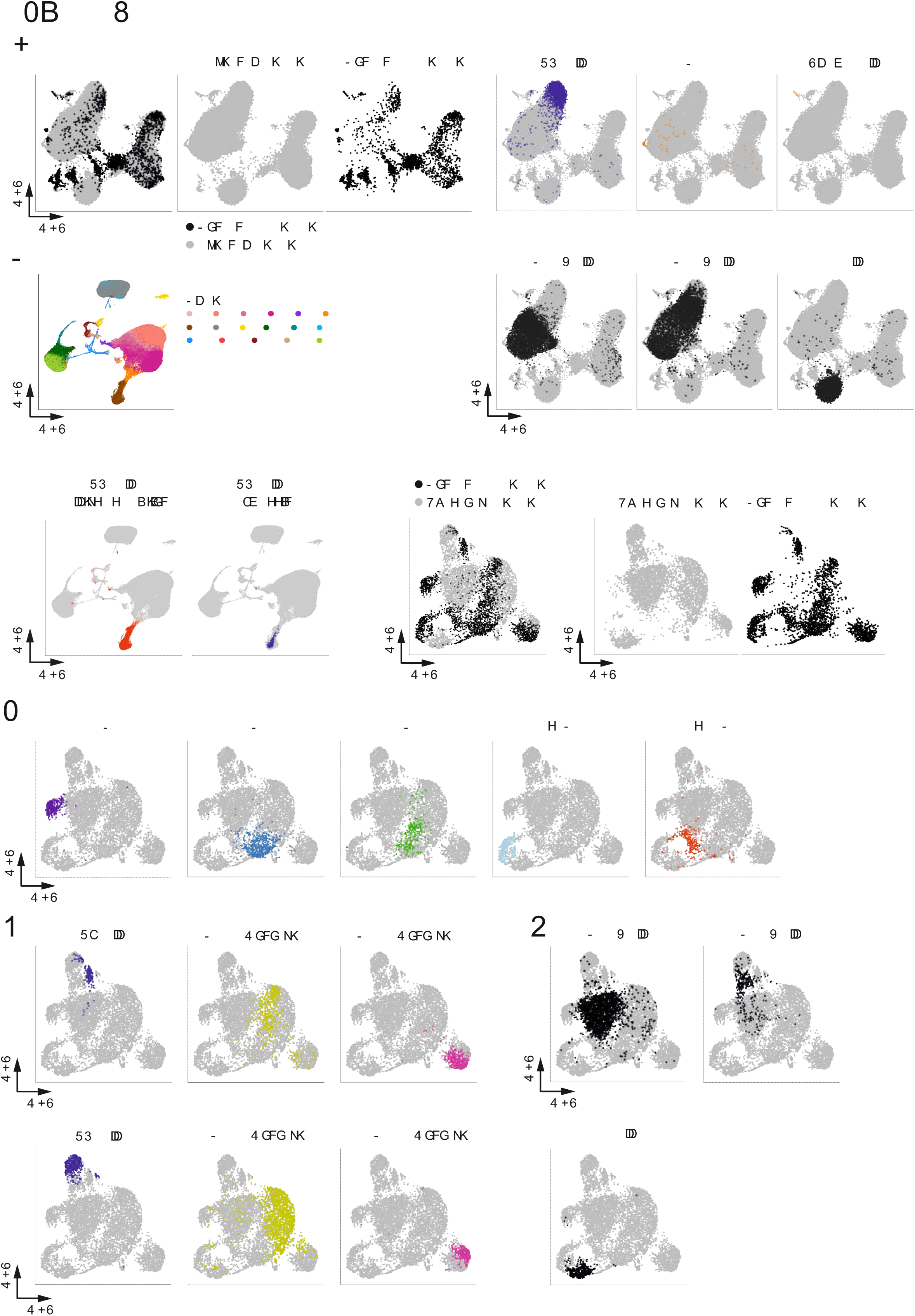
Integration of the new scRNA-seq dataset with three individual PBMC datasets. **(A)** UMAP dimensionality reduction based on the combined dataset of the new mononuclear myeloid scRNA-seq data and the external 33k-PBMC dataset. Cells originating from the new scRNA-seq data are colored black and cells from the external PBMC dataset are colored grey. (**B**) Overlay of cell annotations of cells from NK cells, CD34^+^, plasma cells, CD4^+^ T cells, CD8^+^ T cells, and B cells from the 33-k PBMC dataset. **(C)** Clustering of the HCA dataset. **(D)** Prediction and backmapping of NK cells from the novel scRNA-seq dataset onto the complete HCA dataset. Left UMAP graph shows the computed prediction score for NK cells of the HCA dataset using the new scRNA-seq consensus map information. Red color indicates highest prediction score. Right UMAP graph shows the location of NK cells from the new scRNA-seq data within the combined dataset. **(E)** UMAP dimensionality reduction based on the combined dataset of the new mononuclear myeloid scRNA-seq data-based consensus map and a PBMC dataset processed by the BD Rhapsody technology. Cells originating from the new scRNA-seq consensus map are colored black and cells from the Rhapsody PBMC dataset are colored grey. **(F)** Overlay of cell annotations of the DC subsets from the new scRNA-seq data-based consensus map on the combined UMAP. **(G)** Overlay of cell annotations of NK cells and monocyte subsets identified in the new scRNA-seq data-based consensus map (top panel) and the respective cell annotations of NK cells, CD14^+^ and CD16^+^ monocytes from the Rhapsody PBMC dataset. **(H)** Overlay of cell annotations of CD4^+^ T cells, CD8^+^ T cells and B cells from the external PBMC dataset.

## Table legends

**Tables S1.**
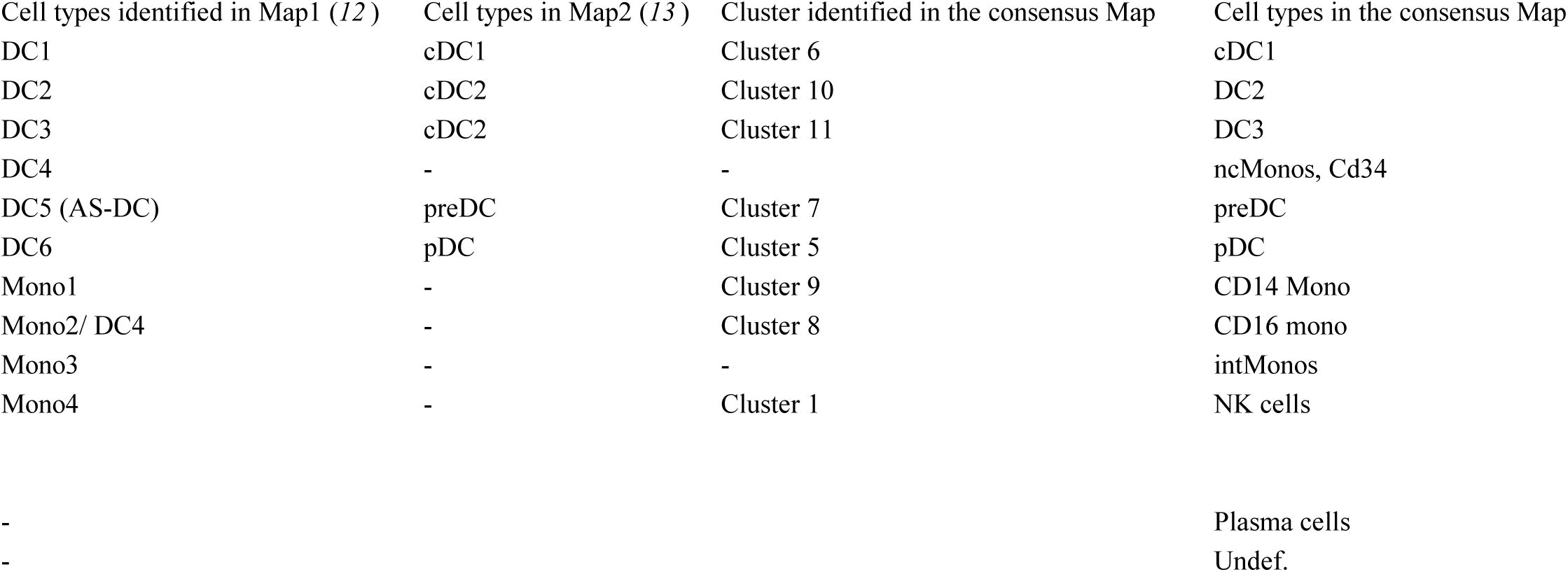
Cell types classified in the respective studies

**Data Table S1:** Data Table S1.csv. Gene signatures of the 11 clusters identified in our new scRNA-seq consensus map

**Data Table S2:** Data Table S2.xlsx. Gene signatures derived from map 2

## EXPERIMENTAL PROCEDURE

### CONTACT FOR REAGENT AND RESOURCE SHARING

Further information and requests for resources and reagents should be directed to and will be fulfilled by the Lead Contact

### EXPERIMENTAL MODEL AND SUBJECT DETAILS

#### Peripheral blood mononuclear cells (PBMC)

Buffy coats or venipuncture blood were obtained from healthy donors (University hospital Bonn, local ethics vote 203/09) after written consent was given according to the Declaration of Helsinki. Peripheral blood mononuclear cells (PBMC) were isolated by Pancoll (PAN-Biotech) density centrifugation from buffy coats.

### METHOD DETAILS

#### Flow cytometric analysis

Whole blood or buffy coat was diluted in room temperature PBS (1:2 or 1:5, respectively) and layered onto polysuccrose solution (Pancoll; PAN Biotech, Germany) for the enrichment of mononuclear cells by density gradient centrifugation according to the manufacturer’s instructions. After three times washing in cold PBS, cells were resuspended in FACS-buffer (0.5% BSA, 2 mM EDTA in PBS) for surface marker staining (Table S2). Human FcR-Block (Miltenyi Biotec, Germany) was included to reduce unspecific staining. After 1 h incubation at 4° in the dark, cells were washed and optionally stained for additional 20 min with 1:400 anti-biotin BV421 in FACS-buffer for CADM1-biotin secondary staining. Washed cells were incubated with L/D Marker DRAQ7 (BioLegend, USA) for 5 min at room temperature before acquisition and sorting of the cells using a BD FACSARIA III (BD BioSciences, USA). Single antibody staining was prepared in parallel to assess fluorescence spillover. Fluorescence-minus-one (FMO) controls were prepared in addition for critical markers to set sorting gates. Post-sort data analysis was performed using FlowJo software (FlowJo, Tree Star Inc., USA). The packages “flowCore” and “flowWorkspace” were used to import raw data into R. For dimensionality reduction with UMAP fluorescence parameters were transformed with logicleTransform (Becht et al., 2018).

#### Library preparation and sequencing using Smart-Seq2

Our new index-sorted single cell transcriptome dataset was based on the Smart-Seq2 protocol (Picelli et al., 2013). For single cell sorting into 384-well plates and to ensure sufficient cell numbers and balanced representation from each main myeloid subset, loose sorting gates have been set covering the entire space of alive CD45^+^Lin (CD3, CD19, CD20, CD56)^-^CD14^+^, CD16^+^ or CD14^-^CD16^-^HLA-DR^+^ cells. To achieve this, the alive CD45^+^Lin^-^ compartment was divided to sort 24 cells per plate each of CD14^+^CD16^-^, CD14^-^CD16^+^, CD14^+^CD16^+^, HLA-DR^+^CADM1^+^, HLA-DR^+^CADM^-^AXL^+^SIGLEC6^+^, HLA-DR^+^CADM1^-^AXL^-^SIGLEC6^-^CD123^+^CD11c^-^, HLA-DR^+^CADM1^-^ AXL^-^SIGLEC6^-^CD123^-^CD11c^-^ or HLA-DR^+^CADM^-^AXL^-^SIGLEC6^-^CD123^-^CD11c^-^ cells. Cells were FACS sorted into eight 384-well plates containing 2.3μl lysis buffer (Guanidine Hydrochloride (50 mM), dNTPs (17.4mM), 2.2μM SMART dT30VN primer) retaining protein expression information for every well to subsequently match with the respective single-cell transcriptomic data in an index sorting approach. Plates were sealed and stored at −80°C until further processing. Smart-Seq2 libraries were finally generated on a Tecan Freedom EVO and Nanodrop II (BioNex) system as previously described (Picelli et al., 2013).

In short, lysed cells were incubated at 95°C for 3 min. 2.7 μl RT mix containing SuperScript II buffer (Invitrogen), 9.3mM DTT, 370mM Betaine, 15mM MgCl2, 9.3U SuperScript II RT (Invitrogen), 1.85U recombinant RNase Inhibitor (Takara), 1.85 μM template-switching oligo was aliquoted to each lysed cell using a Nanodrop II liquid handling system (BioNex) and incubating at 42°C for 90 min and at 70°C for 15min. 7.5μl preamplification mix containing KAPA HiFi HotStart ReadyMix and 2μM ISPCR primers was added to each well and full-length cDNA was amplified for 16 cycles. cDNA was purified with 1X Agencourt AMPure XP beads (Beckman Coulter) and eluted in 14μl nuclease-free water. Concentration and cDNA size were checked for select representative wells using a High Sensitivity DNA5000 assay for the Tapestation 4200 (Agilent). cDNA was diluted to an average of 200pg/μl and 100pg cDNA from each cell was tagmented by adding 1μl TD and 0.5μl ATM from a Nextera XT DNA Library Preparation Kit (Illumina) to 0.5μl diluted cDNA in each well of a fresh 384-well plate. The tagmentation reaction was incubated at 55°C for 8min before removing the Tn5 from the DNA by adding 0.5μl NT buffer per well. 1μl well-specific indexing primer mix from Nextera XT Index Kit v2 Sets A-D and 1.5μl NPM was added to each well and the tagmented cDNA was amplified for 14 cycles according to manufacturer’s specifications. PCR products from all wells were pooled and purified with 1X Agencourt AMPure XP beads (Beckman Coulter) according to manufacturer’s protocol. The fragment size distribution was determined using a High Sensitivity DNA5000 assay for the Tapestation 4200 (Agilent) and library concentration was determined using a Qubit dsDNA HS assay (Thermo Fischer). Libraries were clustered at 1.4pM concentration using High Output v2 chemistry and sequenced on a NextSeq500 system SR 75bp with 2*8bp index reads. Single-cell data was demultiplexed using bcl2fastq2 v2.20.

#### Proteomic Data

To validate the gene signature associated with the mono 4 subset as described by Villani et al. on the protein level, we extracted copy numbers from key signature proteins from the publicly accessible proteomic resource (http://www.immprot.org/) described by Rieckmann et al. containing quantitative high-resolution mass-spectrometry data derived from FACS-enriched human primary blood cells (Rieckmann et al., 2017). Copy numbers were visualized as a heatmap using the *pheatmap* package (v1.0.10) in R.

#### Cytospin preparation and May-Grünwald/Giemsa staining

Cell populations of interest were sorted into 1.5 ml reaction tubes containing 200 μl FACS-buffer using a BD FACSARIA III (BD BioSciences, USA). The cell suspension was centrifuged onto SuperFrost Plus glass slides (Thermo Scientific, USA) at 1000 rpm for 5 min using a Universal 16A slide centrifuge (Andreas Hettich GmbH & Co.KG, Germany). Slides were air-dried overnight and subsequently stained with May-Grünwald/Giesma solution (Carl Roth GmbH, Germany) according to the manufacturer’s guidelines. Images were acquired with a BZ-9000 (Keyence, Japan).

#### Targeted sequencing of human PBMC with the BD Rhapsody™ system

Whole blood was diluted in room temperature PBS (1:2) and layered onto polysuccrose solution (Pancoll; PAN Biotech, Germany) for the enrichment of mononuclear cells by density gradient centrifugation according to the manufacturer’s instructions. Granulocytes were isolated using erythrocyte lysis buffer (ELB, 0.15M NH4Cl, 0.01M KHCO3, 0.1mM EDTA, pH 7.4 at ca. 2-8°C) and mononuclear cells and granulocytes are mixed in a ratio of 2:1. After washing in cold PBS 10.000 cells were loaded onto a BD Rhapsody™ cartridge and processed according to manufacturer’s instructions for targeted single-cell RNA-seq using the predesigned Immune Response Panel (Human). The library was clustered at 1.75pM on a NextSeq500 system (Illumina) to generate ∼40.000 paired end (2*75bp) reads per cell using High Output v2 chemistry. Sequenced single-cell data was demultiplexed using bcl2fastq2 v2.20.

#### Single-cell RNA-Seq raw data processing

Following sequencing by the Smart-Seq2 method (Picelli et al., 2013), RNA-Seq libraries were subjected to initial quality control using FASTQC (http://www.bioinformatics.babraham.ac.uk/projects/fastqc, v0.11.7) implemented in a scRNA pre-processing pipeline (docker image and scripts available at https://hub.docker.com/r/pwlb/rna-seq-pipeline-base/, v0.1.1; https://bitbucket.org/limes_bonn/bulk-rna-kallisto-qc/src/master/, v0.2.1). Next, raw reads were pseudoaligned to the human transcriptome (GRCh38, Gencode v27 primary assembly) using Kallisto with default settings (v0.44.0) (Bray et al., 2016). Based on the pseudoalignment estimated by Kallisto, transcript levels were quantified as transcripts per million reads (TPM). TPM counts were imported into R using tximport (Soneson et al., 2015) and transcript information was summarized on gene-level. We imported the resulting dataset of 43,612 features across 3,072 samples and performed the downstream analysis using the R package Seurat (v.2.3.4, (Butler et al., 2018)).

For processing of the single-cell data obtained by the BD Rhapsody™ system, we run the recommended BD Rhapsody™ Analysis Pipeline of Seven Bridges Genomics (sbgenomics.com/bdgenomics) with standard settings. The resulting count table that was accounted for UMI sequencing and amplification errors, was comprised of 488 features across 7,873 cells. Normalization and further downstream analysis were conducted in R using Seurat (v.2.3.4, (Butler et al., 2018)).

#### Quality control

Concerning our new index-sorted and Smart-Seq2-based single cell transcriptome dataset the following quality control scheme using various meta information was performed to obtain high-quality transcriptome data: 1) We removed genes that are detected in less than 6 cells (0.2 percent of cells), 2) and removed cells that have less than 1,000 uniquely detected genes. Next, we filtered further outlier cells with 3) less than 50,000 unique reads, 4) less than 30% pseudoalignment of reads to the transcriptome, 5) a lower rate of endogenous-to-mitochondrial count rate of 2, 6). This quality control scheme results in a dataset of 29,240 genes across 2,509 cells.

#### Normalization of single-cell transcriptomic data

To reduce the influence of variation of sequencing depth among samples we applied a log-normalization to the data and scaled each cells gene expression profile to a total count of 10,000. In addition, we corrected for other technical effects including differences in the fraction of mitochondrial counts as well of unique detected genes using a linear regression model for these factors. The residuals of this regression are scaled and centered and used for further downstream analysis.

#### Dimensionality reduction and clustering

In order to reduce the dimensionality of the dataset, we selected highly variable genes as genes with an average expression of at least 0.0125 and a scaled dispersion of at least 1. This resulted in a total of 2491 genes, which were used as input for a principal component (PC) analysis. We visualized the standard deviation of the first 20 PCs and identified the first 10 principal components with a minimum standard deviation of at least 2 as significant PCs. Next, we utilized Uniform Manifold Approximation and Projection (UMAP) to further reduce the data into a two-dimensional representation (Becht et al., 2018). To test for cellular heterogeneity, we used a shared nearest neighbor (SNN)-graph based clustering algorithm implemented in the Seurat package. We used the first 10 principal components for constructing the SNN-graph and set the resolution to 1. Monocle was used to infer differentiation trajectories by using the Louvain clustering method, umap dimensionality reduction and the SimplePPT algorithm (Qiu et al., 2017)

#### Additional analysis

Differentially expressed (DE) genes were defined using a Wilcoxon-based test for differential gene expression built in the Seurat pipeline (v.2.3.4) (Data Table S1). Unless otherwise stated genes have been considered as differentially expressed, if the adjusted p-value is smaller than 0.1. Top10 DE genes have been visualized using heatmap of hierarchical clustered gene expression profiles. DE genes have been verified with current literature.

#### Gene signature enrichment analysis

Single-cell RNA-Seq data is inherently sparse and a high-dropout rate is limiting the use of single marker genes to identify cell populations. In order to unambiguously identify the different cell types, we have used an updated version of a gene signature score analysis described earlier (Mass et al., 2016). A cell population is always characterized by genes that are significantly upregulated in comparison to other populations and genes that show significantly lower expression in comparison to the background populations. In order to increase the power, we use both up and downregulated gene signatures for the calculation of the gene expression scores. A cell *i* may be described by a gene expression profile *A*[*i,j*] as the combination of gene expression values of all genes *j*. To calculate a signature score for a cell *i*, we first calculate the scaled average expression of all genes *j_up_* from an upregulated list and of all genes *j_down_* from a downregulated gene list. The difference between these two is scaled and visualized. The visualization is performed as color-coded overlay on the UMAP dimensionality reduction or as density distribution.

#### Data analysis of external single-cell RNA-Seq datasets

To assess the single-cell RNA-Seq data of human dendritic cells and monocytes publicly available under the Gene Expression Omnibus accession number GSE94820, we applied the processing steps previously described (Villani et al., 2017). We focused on the “discovery” dataset and performed downstream analysis with the R software package Seurat (https://github.com/satijalab/seurat; http://satijalab.org/seurat/, v.2.3.4). Uniform manifold approximation (UMAP) algorithm integrated into the Seurat package was used as a dimensionality reduction method with standard settings. To define cell-type specific gene signatures for all cell populations, a Wilcoxon-based test was used. We considered genes as differentially expressed with an adjusted p-value of smaller 0.1 and a log2-fold change of higher than 1 or lower than −1, respectively. A global comparison of all cell types was performed by calculating the Pearson Correlation coefficients between the average expression profiles of all clusters. Scaled gene expression profiles have been used.

In order to have a comprehensive single-cell RNA-Seq dataset of human PBMCs, we downloaded a dataset containing transcriptome data of 33,148 PBMCs from a healthy donor (short 33k-PBMC dataset), which is publicly available on the 10x Genomics webpage (https://support.10xgenomics.com/single-cell-gene-expression/datasets/1.1.0/pbmc33k). Next, we followed the general data analysis scheme described at the Seurat package webpage (https://satijalab.org/seurat/get_started_v1_4.html). Briefly, we used the filtered cell-gene matrix provided by 10x Genomics and imported the data and performed the analysis with the Seurat package. We filtered genes that are expressed in less than three cells and removed cells from the data set that have gene counts for less than 500 genes or for more than 2500 genes. In addition, we removed cells that have more than 5% mitochondrial counts. This resulted in a dataset of 17943 genes across 28.823 cells. Next, a log-normalization was applied, and highly variable genes were identified by applying a dispersion cutoff of 0.8 (2.281 variable genes). To account for technical variability in the dataset, a linear model was used to regress out the effects of the number of measured molecules per cell, the fraction of mitochondrial counts as well as the effect introduced by processing the cells in different sets. The first 25 principal components were used for a graph-based clustering approach. NK cell specific genes were identified by a Wilcoxon-based test for differential gene expression (adj. p-value < smaller 0.1, |log2-fold change| > 1).

#### Backmapping

In order to compare the transcriptome profiles of monocytes isolated from the dataset derived from GSE94820 (Villani et al., 2017) with the comprehensive PBMC dataset, we used the previously introduced canonical correlation alignment to combine datasets (Butler et al., 2018). First, we isolated all monocyte populations from Villani et al. and all monocyte and NK cell populations of the 10x Genomics dataset. Both datasets are normalized, scaled and a linear regression was performed to account for differences in the number of detected genes. In both datasets, a feature selection was performed to identify genes with high dispersion. We determined the mutual highly variable genes as the overlap of the 4.000 genes from each dataset with highest dispersion. Next, we combined both datasets by performing the canonical correlation alignment, which resulted in an integrated dataset comprising 41.620 genes across 8.846 cells. UMAP dimensionality reduction was applied to the dataset using the first 8 canonical correlation alignment components and 40 neighbor points as well as a minimal distance of 0.01.

In addition, we downloaded from the data portal (https://preview.data.humancellatlas.org/) of the HCA consortium a single-cell dataset comprised of immune cells from human cord blood samples. When analyzing this dataset, we observed a donor dependent batch effect and thus decided to use an “anchoring” approach to harmonize the different batches of the single-cell dataset and to integrate the new consensus map. To this end, we took advantage of the R package Seurat (v. 3.0.0.9000). After filtering genes that were expressed in less than 10 cells of the HCA dataset with a cell being kept when 500 genes were detected, we ended up with a large dataset that contained 21,409 genes expressed across 254,937 cells. Next, we merged this Seurat object with the Seurat object of the new consensus map. We treated the different batches of the HCA dataset as individual datasets and normalized them and the expression table of the consensus map separately. For each dataset, we calculated the top 2,000 most variable genes based on a variance stabilizing transformation followed by data integration by leaving the standard settings unaltered. The integrated dataset was visualized using UMAP based on the top 30 computed PCs. For cell type prediction of the cord blood cells based on the calculated clusters of the new consensus map, we followed the recommendations of the Seurat vignette for the ‘FindTransferAnchors’ and the ‘TransferData’ functions. First, we repeated the steps above but without integration of the new consensus map data. We used the resulting integrated HCA dataset as query dataset and the new consensus map as reference dataset. Because of the large cell number of the HCA dataset, we projected the PCA from the query dataset onto the reference dataset. The remaining standard settings were left unaltered. Finally, we transferred the cluster information of the new consensus map onto the query dataset. The resulting prediction scores were visualized as color code onto the UMAP graph by coloring the highest prediction score red. Clustering of the dataset was done based on the construction of an SNN-graph by setting the resolution to 0.6. Cluster 5, 7, 9 and 13 were found to be associated with monocytes or DCs and thus the HCA dataset was filtered on these cells followed by repetition of the abovementioned steps.

#### Population-based gene signatures of pDC, pre-DC, cDC1 and cDC2

Specific gene signatures of up or downregulated in the comparison of human DC subsets have been identified as described earlier using the publicly available dataset (See et al., 2017) (GEO accession number: GSE80171). Gene signatures are available in supplementary Data Table S2.

#### Data visualization

In general, the ggplot2 package was used to generate figures (Wickham, 2016).

### QUANTIFICATION AND STATISTICAL ANALYSIS

Statistical analysis was performed using the R programming language. Statistical tests used are described in the figure legend or methods part, respectively. Differentially expressed genes have been identified using a Wilcoxon-based test for differential gene expression. If not otherwise stated a significance level of 0.1 was applied to adjusted p-values (Benjamini Hochberg).

### DATA AND SOFTWARE AVAILABILITY

Processed and raw scRNA-seq datasets are available through the Gene Expression Omnibus (GSE126422). Additional Data tables are provided in form of EXCEL Tables (Data S1, S2)

#### Data Table S1: Data Table S1.csv

Gene signatures of the 11 clusters identified in our new scRNA-seq consensus map

#### Data Table S2: Data Table S2.xlsx

Gene signatures derived from map 2

### ADDITIONAL RESOURCES

In addition, we provide an interactive web tool to visualize the single-cell RNA-Seq data together with the flow cytometry data at https://paguen.shinyapps.io/DC_MONO/ (external database S1).

