## Supplementary material for "A rule-based data-informed cellular consensus map of the human mononuclear phagocyte cell space": Antibody panel

**Table S2.** **List of antibodies and secondary staining reagents for flow cytometric analysis.**

| **Usage** | **Name** | **Conjugate** | **Clone** | **Source** |
| --- | --- | --- | --- | --- |
| Lineage, Fig.1,2,3A-F,4A-C | CD3 | APC-Cy7 | OKT3 | BioLegend |
| Lineage, Fig.1,2,3A-F,4A-C | CD19 | APC-Cy7 | SJ25C1 | BioLegend |
| Lineage, Fig.1,2,3A-F,4A-C | CD20 | APC-Cy7 | 2H7 | BioLegend |
| Lineage, Fig.1,2,3A-F,4A-C | CD56 | APC-Cy7 | 5.1H11 | BioLegend |
| Fig.1,2,3A-F,4A-C | HLA-DR | BV570 | L243 | BioLegend |
| Fig.1,2,3A-F,4A-C | CD16 | BV510 | 3G8 | BioLegend |
| Fig.1,2,3A-F,4A-C | CD14 | BV711 | M5E2 | BioLegend |
| Fig.1,2,3A-F,4A-C | CD45 | PE/Cy7 | HI30 | BioLegend |
| Fig.1,2,3A-F,4A-C | CD11c | PerCP | Bu15 | BioLegend |
| Fig.1,2,3A-F,4A-C | CD45RA | FITC | HI100 | BioLegend |
| Fig.1,2,3A-F,4A-C | CD34 | AF700 | 581 | BioLegend |
| Fig.1,2,3A-F,4A-C | CD1C | PE/CF594 | L161 | BioLegend |
| Fig.1,2,3A-F,4A-C | SIGLEC6 | PE | 767329 | R&D Systems |
| Fig.1,2,3A-F,4A-C | CD123 | BV785 | 6H6 | BioLegend |
| Fig.1,2,3A-F,4A-C | CADM1 | Biotin | 3E1 | MBL |
| Fig.1,2,3A-F,4A-C | CD33 | BV650 | WM53 | BioLegend |
| Fig.1,2,3A-F,4A-C | AXL | APC | 108724 | R&D Systems |
| Fig.1,2,3A-F,4A-C | Streptavidin | BV421 | - | BioLegend |

| **Usage** | **Name** | **Conjugate** | **Clone** | **Source** |
| --- | --- | --- | --- | --- |
| Lineage, Fig.4D | CD3 | APC-Cy7 | OKT3 | BioLegend |
| Lineage, Fig.4D | CD19 | APC-Cy7 | SJ25C1 | BioLegend |
| Lineage, Fig.4D | CD20 | APC-Cy7 | 2H7 | BioLegend |
| Fig.4D | HLA-DR | BV570 | L243 | BioLegend |
| Fig.4D | CD16 | BV510 | 3G8 | BioLegend |
| Fig.4D | CD14 | BV711 | M5E2 | BioLegend |
| Fig.4D | CD45 | PE/Cy7 | HI30 | BioLegend |
| Fig.4D | CD11c | PerCP | Bu15 | BioLegend |
| Fig.4D | CCR3 | FITC | REA574 | Miltenyi |
| Fig.4D | CD66b | AF700 | G10F5 | Biolegend |
| Fig.4D | CD7 | PE/Dazzle | CD7-6B7 | Biolegend |
| Fig.4D | CD160 | PE | BY55 | Biolegend |
| Fig.4D | CD107a | BV785 | H4A3 clone | Biolegend |
| Fig.4D | NCR1/NKp46 | BV421 | 9E2 | Biolegend |
| Fig.4D | CD56 | BV650 | 5.1H11 | Biolegend |

| **Usage** | **Name** | **Conjugate** | **Clone** | **Source** |
| --- | --- | --- | --- | --- |
| Lineage, Fig.3G-H | CD3 | APC-Cy7 | OKT3 | BioLegend |
| Lineage, Fig.3G-H | CD19 | APC-Cy7 | SJ25C1 | BioLegend |
| Lineage, Fig.3G-H | CD20 | APC-Cy7 | 2H7 | BioLegend |
| Fig.3G-H | HLA-DR | BV570 | L243 | BioLegend |
| Fig.3G-H | CD16 | BV510 | 3G8 | BioLegend |
| Fig.3G-H | CD14 | BV711 | M5E2 | BioLegend |
| Fig.3G-H | CD45 | PE/Cy7 | HI30 | BioLegend |
| Fig.3G-H | CD11c | PerCP | Bu15 | BioLegend |
| Fig.3G-H | CD45RA | FITC | HI100 | BioLegend |
| Fig.3G-H | CD34 | AF700 | 581 | BioLegend |
| Fig.3G-H | CD1C | PE/CF594 | L161 | BioLegend |
| Fig.3G-H | CD123 | BV785 | 6H6 | BioLegend |
| Fig.3G-H | CADM1 | Biotin | 3E1 | MBL |
| Fig.3G-H | Streptavidin | BV421 | - | BioLegend |
| Fig.3G-H | CD56 | BV650 | 5.1H11 | Biolegend |
| Fig.3G-H | Slan | APC | M-DC8 | Miltenyi |
